# Predicting protein curvature sensing across membrane compositions with a bilayer continuum model

**DOI:** 10.1101/2024.01.15.575755

**Authors:** Yiben Fu, David H. Johnson, Andrew H. Beaven, Alexander J. Sodt, Wade F. Zeno, Margaret E. Johnson

## Abstract

Cytoplasmic proteins must recruit to membranes to function in processes such as endocytosis and cell division. Many of these proteins recognize not only the chemical structure of the membrane lipids, but the curvature of the surface, binding more strongly to more highly curved surfaces, or ‘curvature sensing’. Curvature sensing by amphipathic helices is known to vary with membrane bending rigidity, but changes to lipid composition can simultaneously alter membrane thickness, spontaneous curvature, and leaflet symmetry, thus far preventing a systematic characterization of lipid composition on such curvature sensing through either experiment or simulation. Here we develop and apply a bilayer continuum membrane model that can tractably address this gap, quantifying how controlled changes to each material property can favor or disfavor protein curvature sensing. We evaluate both energetic and structural changes to vesicles upon helix insertion, with strong agreement to new *in vitro* experiments and all-atom MD simulations, respectively. Our membrane model builds on previous work to include both monolayers of the bilayer via representation by continuous triangular meshes. We introduce a coupling energy that captures the incompressibility of the membrane and the established energetics of lipid tilt. In agreement with experiment, our model predicts stronger curvature sensing in membranes with distinct tail groups (POPC vs DOPC vs DLPC), despite having identical head-group chemistry; the model shows that the primary driving force for weaker curvature sensing in DLPC is that it is thinner, and more wedge shaped. Somewhat surprisingly, asymmetry in lipid shape composition between the two leaflets has a negligible contribution to membrane mechanics following insertion. Our multi-scale approach can be used to quantitatively and efficiently predict how changes to membrane composition in flat to highly curved surfaces alter membrane energetics driven by proteins, a mechanism that helps proteins target membranes at the correct time and place.

**Significance:** Proteins must recruit to membranes for essential biological functions including endocytosis and cell division. In addition to recognizing specific lipid head-groups, many of these proteins also ‘sense’ the curvature of the membrane, but the strength of sensing is known to vary with distinct membrane compositions. Predicting the dependence of sensing on changes to lipid composition cannot be done *a priori* due to the multiple material properties, including bilayer thickness, bending rigidity, tilt modulus, spontaneous curvature, and leaflet asymmetry that vary with lipid type. Here we use a multi-scale approach to systematically address this gap, developing a double-leaflet continuum model that is informed by structural deformations from all-atom MD and validated against in vitro experiments. This efficient approach can be applied and extended to quantify how proteins sense and drive membrane curvature across a wide range of membrane bilayers, including distinct leaflet compositions and membrane geometries.

## I. Introduction

Cytoplasmic proteins that bind biological membranes are essential for membrane remodeling pathways like clathrin-mediated endocytosis [1, 2] and cell division [3]. Protein domains that mediate membrane binding are often classified by their recognition of specific lipid head groups such as PI(4,5)P_2_ [4-6]. However, experimental studies have shown how multiple membrane binding domains can also recognize membrane curvature[7], including the epsin N-terminal homology (ENTH) domain, the bin-amphiphysin-rvs (BAR) domain [8], and septins[3, 9]. For the ENTH domain, this curvature ‘sensing’ ability stems from its N-terminal amphipathic helix, which inserts into the outer leaflet of the bilayer, generating asymmetry in local lipid packing and tension between the top and bottom leaflet [10, 11]. Helix inserting domains like ENTH bind more strongly to more highly curved vesicles, and truncation of this helix effectively eliminates the curvature sensing ability[10]. These amphipathic, membrane-inserting helices are common to a variety of membrane binding proteins (e.g. septins[12], arfGAP[13], BAR-domain containing proteins[14]), but they do not all exhibit the same curvature sensing abilities across experimental conditions. While this can in part stem from differences in the sequence or length of the helices[15], the composition of the membrane also plays a role, as the insertion of the helix locally perturbs the membrane[16]. Because changing the membrane lipid composition will simultaneously influence multiple material properties of the membrane, including bending rigidity, thickness, lipid tilt, compressibility, and leaflet spontaneous curvature, it has not been possible to systematically quantify how lipid composition impacts curvature sensing. However, many of these material parameters have been measured for distinct lipid types from experiment [17-20] and atomistic simulations [21, 22], including lipid spontaneous curvature which largely reports on the shape of the lipid, shifting positive for wedge-shaped lipids that taper into the midplane (like DLPC) and zero for cylindrical lipids. Here we use a continuum bilayer membrane model that controls for each of these material properties of the membrane to perform a quantitative analysis of curvature sensing by inserting helices into vesicles from highly curved (R=10nm) to flat surfaces. The structure of the membrane around the helix insertion is validated against MD simulations on flat bilayers, and we show that our model predicts curvature sensing in agreement with new *in vitro* experiments using the ENTH domain and previously published *in vitro* measurements [10].

Computation and theory provide highly controlled and tunable methods to quantify the energetic and structural changes arising from helix insertion in membrane bilayers[23]. However, tradeoffs in capturing molecular detail vs micron-scale membrane surfaces have thus far limited a quantitative comparison of lipid composition and average membrane thickness on curvature sensing. At the molecular scale, molecular dynamics (MD) simulations have measured the distribution of membrane stress[24] due to helix insertion [25] and shown how defects in the local ordering of the lipid membrane can regulate helix insertion on flat membranes [26, 27]. On curved membranes, MD simulations characterized how helix insertions can orient along the membrane axis with lower curvature[28], can radially alter local lipid order [29], and can spatially partition based on varying curvature[30]. Computational expense limits MD to the nanometer scale, preventing characterization at a range of curvatures. Classic elasticity theory allows a range of elastic moduli for evaluating solid-like deformations along the membrane thickness/height axis, applicable to nearly arbitrary lengthscales [31] while modeling the large local stresses resulting from helix insertion [32]. Ref. [31] employed an efficient two-dimensional *solid* model, translationally invariant along the third dimension, the length of the cylindrical axis. The two-dimensional solid model requires assuming the helix perturbation extends infinitely along that axis, whereas they are typically only a few nanometers in length. The continuum *surface* models that we use here (e.g., Helfrich-Canham-Evans, or HCE[15, 33]) sit at an intermediate or mesoscale resolution that is computationally much more efficient than MD simulations and more broadly applicable than classical elasticity theory-based solid continuum models. While they lack the single-lipid resolution of molecular approaches, they can simultaneously model energetic and structural changes of membranes at the micron scale in response to a range of perturbations [15, 33, 34], as well as membrane dynamics coupled to protein field variables[35]. By representing the surface in 3D space using points on a 2D triangular mesh, these models can capture realistic membrane shapes with arbitrary topologies [33] as opposed to models that rely on Fourier techniques, which are only capable of modeling small deviations from planarity.

In recent work we implemented such a thin-film continuum surface model that successfully predicted curvature sensing by helix insertion with excellent agreement to experiment, further quantifying how bending rigidity and helix size influenced curvature sensing [15]. However, this single-layer membrane lacks any description of internal membrane structure that is necessary to model the nanometer-scale differences of the perturbations from biological proteins. Furthermore, it offers no clear way to vary the thickness of the bilayer or any properties that couple to thickness. Asymmetry, in particular, is more easily captured in a leaflet-resolved model, compared to a midplane model. As a result, it has not been possible to evaluate how compositional variations affecting the height and tilt moduli, or asymmetry across the two leaflets, alter curvature sensing, despite its significance in biological membranes [36]. For example, the plasma membrane is known to be actively maintained with a highly asymmetric composition, resulting in distinct curvature preferences and bending rigidities for the two leaflets [37]. Further, lipids with different tail groups are measured to have distinct tilt moduli [20], which influences membrane thickness and how it responds to perturbation. Transmembrane proteins have a thickness that influences their partitioning within membranes of complex compositions, preferring to match their thickness to similarly sized bilayers to minimize their energy[38] and control protein conformations[39]. We therefore developed here a multi-layer continuum surface model that addresses these limitations of the single-layer model (Fig 1), capturing distinctions between the two leaflets and the role of lipid height and tilt in membrane structure and energetics.

**Figure 1.**
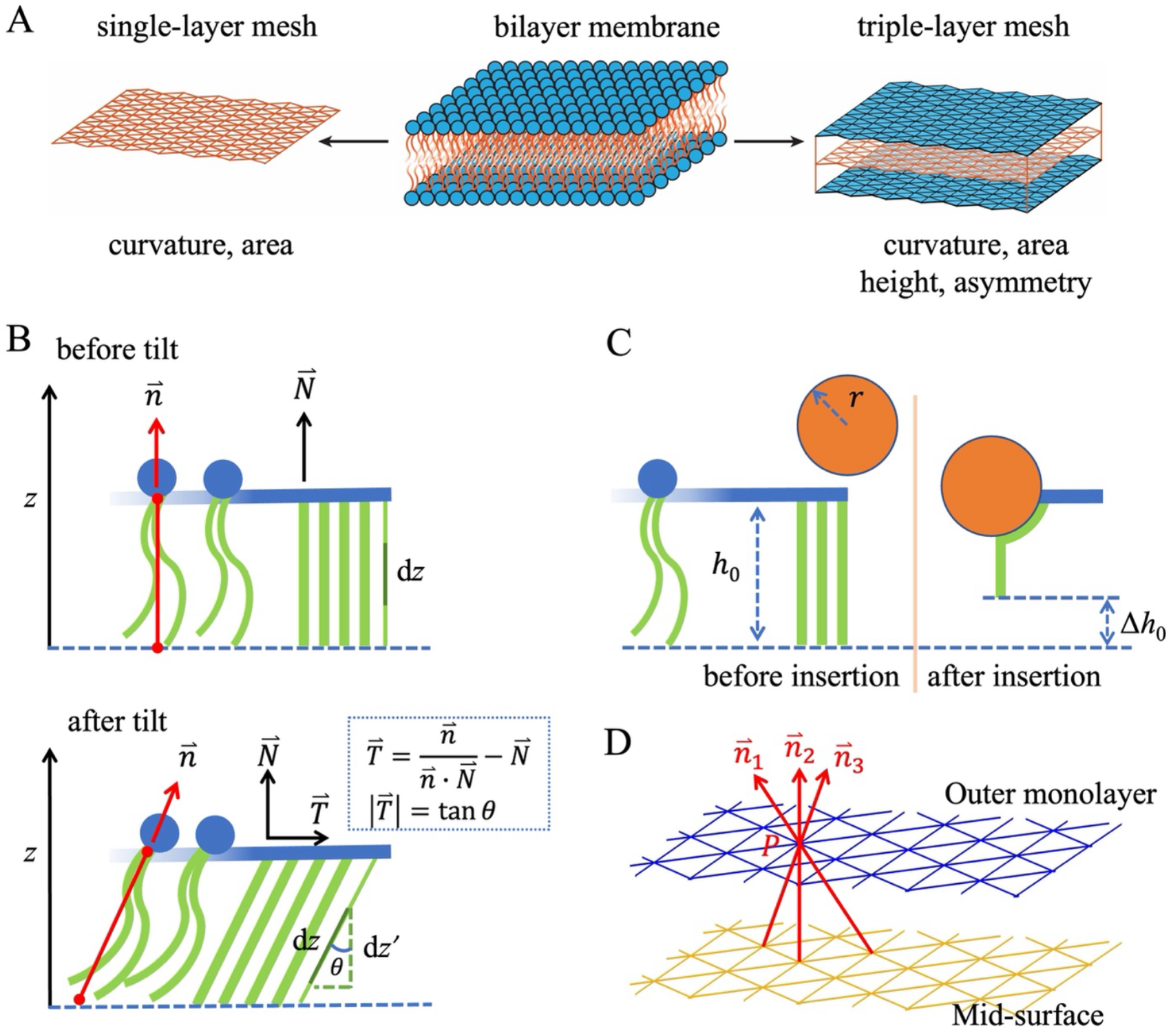
The continuum membrane model is extended here to capture both leaflets of the membrane bilayer coupled via leaflet height and lipid tilt. A) In the middle panel of the top row, we illustrate a bilayer membrane consists of two lipid monolayers, with tail groups in orange and head groups in blue. The left panel shows a continuum membrane model using a single-layer triangular mesh, which represents the mid-surface of the bilayer membrane. This model accurately captures changes in membrane curvature and area elasticity. The right panel shows the new continuum membrane model with a triple-layer of triangular meshes. This new model simulates both monolayers simultaneously, allowing for the analysis of curvature changes and area elasticity in each monolayer, as well as the deformation of the leaflet height arising from curvature changes and lipid tilt. It also supports asymmetry in leaflet material properties. Our model assumes volume incompressibility, such that any changes in area due to curvature result in changes to equilibrium height. (A) Definitions of monolayer normal 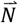, lipid orientation or director 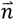, and the lipid tilt 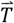, which is a vector parallel to the surface plane with a magnitude 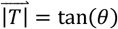. (B) Schematic to show the monolayer height decreases after the α-helix insertion due to lipids undergoing tilt around the insertion. (C) Because the definition of a lipid tilt vector requires a decision on which point P on a layer connects to the other to define the lipid director 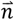, we find reformulating tilt energy via the height difference rather than the tilt vector field is a more straightforward implementation for numerical evaluation of tilt energy. The mesh does not move according to lipid diffusion, and particularly around the insertion, identifying vectors is arbitrary due to the increased number of mesh points on the outer leaflet.

A key development for the multi-layer continuum model is to incorporate the coupling between the two leaflets. Variations between leaflets can be modeled at a range of resolution. The curvature-inducing effect of differential stress[40] on an HCE surface can be captured with the area-difference elasticity (ADE) model [41, 42]. However, the ADE model assumes a uniform membrane thickness throughout the surface [41], and thus cannot model local thickness changes induced by helix insertions. Alternatively, a dual-leaflet Helfrich model with the so-called *Monge* representation can be formulated to describe thickness [38, 43, 44]. However, this formulation is only applicable for relatively small deformations, typically with Fourier analysis; it is not designed for the highly localized deformation underneath the helix.

In our model, we couple the two leaflets via a height variable for both the inner and outer leaflet relative to a midsurface mesh. To define the energy function arising from this variable, we will assume 1) the membrane is incompressible, meaning that lateral stretching of the membrane induces vertical reduction in the thickness, and 2) that lipid tilt can contribute to leaflet thinning. Conceptually, this means that if the curvature of a leaflet changes at any position on the surface, this alters the local stretching of the membrane which will therefore change the equilibrium height of the leaflet at that position. Further deviation from this curvature-corrected equilibrium height can occur due to lipid tilt, which in our model is induced via the helix insertion. We define the energy function based off the tilt-augmented HCE model of Hamm and Kozlov[45], which is derived from classical elasticity theory (see for example Ref [46]). The lipid tilt enters as a vector field, as the thickness is no longer represented as an explicit variable. As we describe in the Methods, we reintroduce thickness explicitly via our leaflet height variables, and we reformulate the tilt energy via the height variable to avoid having to specify lipid tilt by identifying vectors between the leaflets, which is challenging around the helix insertion. We are careful to avoid double-counting of energy from height variations due to curvature vs tilt via our definition of a curvature-dependent equilibrium height. We note that recent work from Terzi and Deserno has identified an additional term in the surface energy functional due to coupling between tilt and curvature gradients[47, 48]. We have neglected this component here as it is parameterized by a poorly known modulus (Gaussian curvature), and it is not readily calculated in our model without a vectoral definition of tilt. The key element that our model is designed to capture is the local perturbation to the lipid structure around a helix insertion, which causes lipid tilt and apparent membrane thinning.

In this paper, we provide our derivation of the coupled two-leaflet energy function based off of the area-difference-elasticity model [41, 42] and the tilt-augmented Helfrich model by Hamm and Kozlov [45]. Our numerical implementation of the continuum membrane bilayer builds from our one-layer surface [15, 49] to introduce an outer mesh layer, an inner mesh layer, and the midsurface layer between the two monolayers, allowing for variations in the local leaflet height/tilt and material properties of each leaflet while maintaining coupling between the leaflets via the height. We performed MD simulations to characterize the local height change introduced to the membrane by the helix insertion, calibrating our continuum model. We performed *in vitro* experiments on vesicles with the same lipid head-group chemistry but varying tail groups, contrasting DOPC, DLPC, and POPC. Using literature values for the material properties of these membranes, our model can predict how the strength of curvature sensing changes with lipid composition, in excellent agreement with the experiments. Our results provide a foundation for predicting how distinct membranes will impact protein recruitment as key material properties vary.

## II. Models and Methods

### II.A Theory of the continuum bilayer Hamiltonian

#### II.A.1. Background on theoretical expression for bilayer membrane energy

Lipid bilayers are composed of two lipid monolayers or leaflets in contact via their hydrophobic tails. The energy of the bilayer membrane can be defined by the sum of the individual mechanical energies of the each monolayers, an energy due to the geometrical constraint between the two monolayers [41], and any external work on the system.

We thus define the total energy of the system as:

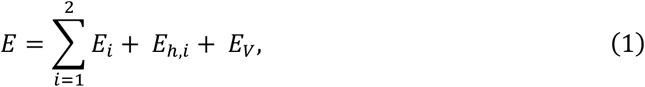

where *E*_*i*_ is the monolayer surface energy, and the index *i* = 1, 2 refers to the inner and outer monolayer, respectively. *E*_*h,i*_ is the energy due to height variations of each leaflet, which is the only term coupling together the two surface layers (discussed below). For the enclosed membrane, like the vesicle membrane, the energy term *E*_*V*_ is incorporated to represent the restriction on the volume enclosed by the membrane,

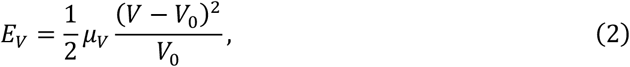

where *μ*_*V*_ is the coefficient of the volume constraint, and *V*_0_ is the equilibrium or target volume. The volume constraint is an equivalent description of the difference between the osmotic pressures inside and outside the vesicle [50]. Previous work found the volume constraint contributes minimally to the overall energetics given the stable difference between the osmotic pressure inside and outside the vesicle, even when perturbed by *α*-helix insertion [15]. For unenclosed or infinite membranes, such as cylindrical and planar membranes, there is no volume constraint or *E*_*V*_ term. The mechanical energy of the *i*th monolayer is given as the HCE Hamiltonian [41, 51, 52],

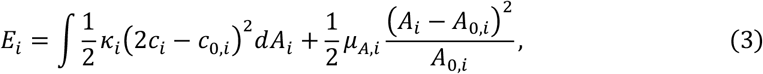

where the parameters are the bending modulus *κ*_*i*_, the spontaneous curvature *c*_0,*i*_, the area elasticity modulus *μ*_*A,i*_, and the equilibrium area *A*_0,*i*_. The energy varies due to changes to the mean curvature *c*_*i*_, which is evaluated at each point on the surface, and the surface area *A*_*i*_. For the derivations in the next sections (not the numerical simulations) we will make the simplifying assumption that the monolayers have the same composition and thus physical properties, *i*.*e*. *κ*_1 =_ *κ*_2_, *μ*_*A*,1 =_ *μ*_*A*,2_, and *c*_0,1_ = −*c*_0,2_ where the minus represents that the two monolayers have opposite lipid orientation. Constants *κ*, *μ*_*A*_, *c*_0_ represent the single leaflet values (monolayer bending modulus, area elasticity modulus and spontaneous curvature, respectively).

#### II.A.2. Derivation of the monolayer equilibrium area

Values of parameters in Eq (3), namely *κ*, *μ*_*A*_ and c_0_, rely on the lipid composition and can be obtained from prior experimental data. For the parameter *A*_0,*i*_, however, its value is dependent on these other parameters and the geometry of the membrane, and we define this relationship here. For a curved membrane such as a sphere, the inner and outer leaflet radii are distinct given a membrane height *h>0*. The area of each leaflet will adjust to relax the membrane compression on the inner layer and stretching on the outer layer (via, e.g. lipid flipping [53]), where we assume the mid-surface area *A*_*m*_ and the membrane height *h* remain fixed. The equilibrium area of each monolayer can be defined as:

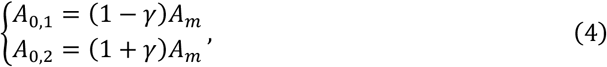

where γ quantifies the relative area change from *A*_*m*_. Note the sum of the equilibrium areas of the two monolayers is independent of γ, *A*_0,1_ + *A*_0,2_ = 2*A*_*m*_, preserving mass conservation. For a flat membrane without external force, its two monolayers will have equal areas, *A*_0,1_ = *A*_0,2_ = *A*_*m*_, such that *γ*_flat_ = 0. The exact value of *γ* depends on the geometry of the surface but can be derived by minimizing the corresponding membrane energy relative to it, 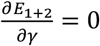. We perform this derivation in the SI and use those exact expressions in our numerical evaluations (SI). For intuition, we can show that for the sphere, when *R* is large (SI):

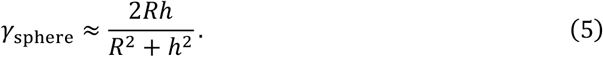

As expected, with flatter membranes, *R* → +∞, and *γ*_sphere_ →0. The difference between the two monolayers’ equilibrium areas in this limit is Δ*A*_0_ = *A*_0,2_– *A*_0,1_ = 16*πR*^3^*h*/(*R*^2^+ *h*^2^). For the large radius *R* ≫ *h*, we have Δ*A*_0_ = 16*πRh*, which is the same as the prediction of the ADE model [42].

#### II.A.3. Background on membrane surface energy with curvature and lipid tilt

Previous work from Hamm and Kozlov[45] derived a membrane surface energy starting from classical elasticity theory that contains monolayer energy terms due to both curvature and lipid tilt. The derivation integrates over the membrane thickness dimension, and results in definitions of the bending modulus, area/thickness modulus, and tilt modulus that are fully constrained by the local elastic constants of the three-dimensional structure [45]. The membrane is assumed incompressible, and no explicit height or thickness variable is present because it has been integrated out. We repeat here key elements that are helpful for our subsequent section. The energy per unit surface area (in contrast to the total energy of Eq. 3) is:

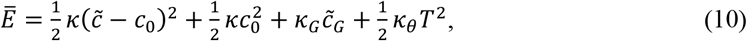

where the additional parameters are the tilt modulus *κ*_*㮈*_, and the Gaussian curvature modulus *κ*_*G*_, and the variables are the effective curvature 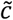, the effective Gaussian curvature 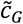, and the tilt magnitude *T*^2^. The effective curvature and effective Gaussian curvature equal their standard definitions when the tilt tensor vanishes [45]. The tilt is a vector measuring the difference between the surface normal 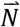 and the local mean lipid director 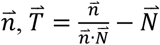, Fig 1. The tilt vector is thus parallel to the midsurface, 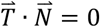.

The surface energy of Eq. 10 derives from a 3D elastic solid representation where the tilt captures the shear strain at the midplane. The area strain at the midplane is given by ε = (*dx*′*dy*′ − *dxdy*)/*dxdy* and the thickness strain at the midplane is given by *ρ* = (*dz*′ − *dz*)/*dz*, where the prime indicates the size of the base of a solid element after deformation, and nonprime is before deformation. The thickness of the leaflet is constrained by an assumption of volume incompressibility that can be stated as:

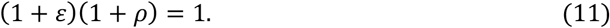

The relative area expansion is related to the membrane curvature via [45, 47],

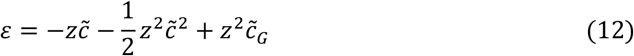

where here we neglect the extra term due to tilt coupled to the curvature gradient [47].

Because this energy function applies to a 2D surface with a tilt vector field, we will in the next section define an approximate energy function that explicitly depends on leaflet height and retains a contribution from lipid tilt.

#### II.A.4. Derivation of a height energy function that accounts for volume incompressibility and tilt

Thickness specifically refers to the (possibly strained) length of the lipid, whereas height refers to the position of the surface above the midplane. In our derivation, we infer variations of the tilt from the height of the leaflets, assuming that deformations in the leaflet induced by the helix are purely tilt, rather than thickness. The tilt energy is thus cast in terms of the height of the leaflet as opposed to an explicit tilt variable. With the introduction here of an explicit height variable *h* in addition to the pre-existing curvature and area surface energy, the height changes cannot be fully independent of curvature variation due to the volume incompressibility of the membrane. The monolayers have an equilibrium height parameter *h*_0_ defined by the lipid hydrocarbon tail length in a flat membrane. Now if the membrane curves, the area and thus the height change to maintain volume incompressibility, and therefore we define a curvature-modified equilibrium height:

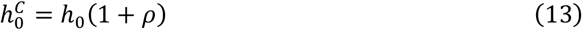

Or equivalently from Eq. 11: 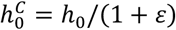. So our values of 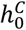 will be defined based on the local curvature ε from Eq. 12 by setting *z* = *h*_0_.

We next define an apparent height strain that accounts for curvature thinning/thickening as

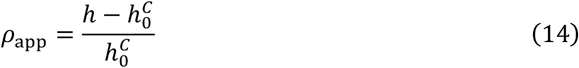

where *h* is the instantaneous height variable. This now reports on energy strain due to additional height changes that are not attributable to curvature or area changes, and are therefore arising from tilt or shear stress.

The fourth term of the energy in Eq. 10, ^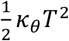^, arises from tilt, and here we will redefine it via our explicit height variable, by relating *T*^2^and *ρ*_app_. Deformations of tilt and area are assumed to be small, and thus the lipid tilt can be related to the relative curvature-modified height change, 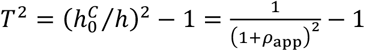. The variable *ρ*_app_ is only physically meaningful as defined over the domain −1 < *ρ*_app_ ≤ 0 or 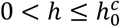, because lipid tilt can only lead to thinning or height decrease of the membrane, and not thickening. If we Taylor expand and keep terms to order 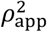 which will be accurate in a limit of small relative deformations (*ρ* ≪ 1), then

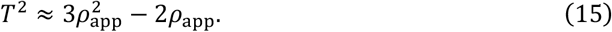

The total energy of the monolayer height deformation is obtained by the integral of the tilt energy term along the whole surface,

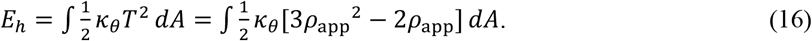

If we define a height modulus *κ*_*h*_ = 3*κ*_㮈_, and complete the square in Eq. 16, we can write the monolayer height energy as

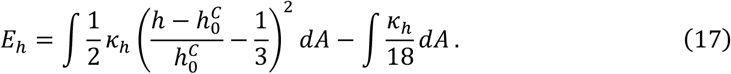

The second term in this equation is a constant, so when measuring energy differences it will cancel out. The bilayer height energy will be the sum of contributions from Eq. 17 for each monolayer. Note that 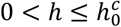, as tilt can only thin the membrane. However, Eq. 17 indicates that the height/tilt energy is minimized when <u>^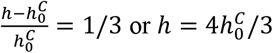^</u>. That is when the observed height is in fact bigger than the curvature-modified equilibrium height. If we add the additional constraint that 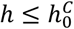, then again the energy will be minimized when 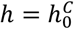. We thus use another term to the Energy per surface area (Eq. 10) to prevent positive values of *ρ*_app_, see SI for details.

#### II.A.5. Response of the membrane energy function due to helix insertion

When an α-helix inserts into the outer monolayer surface of the membrane, it introduces perturbations to the membrane structure and subsequently affects the membrane energy. To account for the influence of helix insertion, we modify the membrane energy at three key points based on the outer monolayer surface patch that is designated as the helix insertion site.

i. Area: The helix insertion reaches to the hydrophobic regions of the outer monolayer, so the neutral surface area of the outer monolayer needs to include the area occupied by the helix. In our model, the effective equilibrium area of the outer monolayer is changed to *A*_0,2_+ *a*, where *a* is the helix insertion area. For the N-terminal α-helix of ENTH protein, *a* is about 2 nm^2^ with 1 nm in width and 2 nm in length. *A*_0,1_ remains unchanged since the helix insertion is not assumed to induce lipid flip-flop.
ii. Tilt/Height: When the helix inserts into the outer monolayer, it causes the adjacent lipids to deform and wrap around the helix, as shown in the MD simulations. As a result, the membrane appears thinner at the insertion site (as depicted in Fig 1), which we will parameterize by a decrease in membrane height Δ*h*_0_ at the insertion patch. For intuition, if one helix with *r* = 0.5 nm and half immersed (as shown in Fig 1), we calculate Δ*h*_0_ to be approximately 0.29 nm, while for completely immersed, Δ*h*_0_ is around 0.57 nm. Deeper insertions thus cause more thinning of the membrane. In the following, we calibrate Δ*h*_0_ to reproduce what is observed in the MD simulations. At the insertion patch, we thus will set the equilibrium height to be 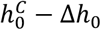. Importantly, although the parameter Δ*h*_0_ defines the monolayer equilibrium height at the insertion, the observed decrease of the monolayer height *h* following energy minimization is typically less due to competing energy terms.
iii. Spontaneous curvature: The spontaneous curvature of the helix insertion, denoted as *c*_0,*ins*_, differs from the spontaneous curvature of the lipid monolayer *c*_0_. We specificy *c*_0,*ins*_ at the insertion patch, while the rest of the surface uses *c*_0_. The inner monolayer spontaneous curvature is not affected by the insertion to the outer layer.

#### II.A.6. Comparing membrane energies to helix binding affinities

Our model measures changes in membrane energies (and membrane shape) in response to perturbations. In this paper, the perturbation is the insertion of an amphipathic helix into the outer leaflet of a spherical membrane. To compare directly with experiment, we follow the same approach as in our previous work[15], connecting the membrane energy to the experimentally measured equilibrium binding affinity of the helix to the surface as it changes with curvature. Our experiments observe wild-type ENTH proteins recruited to small unilamellar vesicles (SUVs) of different radii. By recording the equilibrium protein density on the vesicle *ρ_EL_^eq^*, and with knowledge of the PIP_2_ density*ρ_L_^eq^* and the solution concentration of the protein *ρ_E_^eq^*, we can calculate the dissociation constant of ENTH for different vesicles, *K_D_* = *ρ_E_^eq^ρ_L_^eq^*/*ρ_EL_^eq^*. Both lipid (PIP_2_) and solution protein density are in large excess of the bound protein-lipid complex, meaning *ρ_E_^eq^* → *ρ_E_^tot^* and *ρ_L_^eq^* → *ρ_L_^tot^*. The experimental value, dependent on curvature, is then 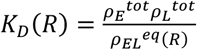. The well-known relation connects to the binding free energy via 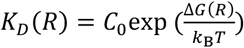 where *C*_0_ = 1*M, k*_B_ is Boltzmann’s constant, and *T* is the temperature. The free energy difference is between 1) the unbound helix and the unperturbed membrane, vs 2) the bound helix to the bound and perturbed membrane which we define with components Δ*G*(*R*) = −*μ* + ε + Δ*E*(*R*), where *μ* is the chemical potential of the helix interaction due to electrostatics, ε is a contribution due to possible conformational changes, and Δ*E*(*R*) is a mechanical energy due to the helix perturbation to the membrane. As we did previously, we assume only the mechanical contribution due to the membrane energy change is curvature dependent, as the ENTH domain without helix insertion binds to membranes without any curvature dependence[15], such that −*μ* + ε is constant. Rearranging, we recover that

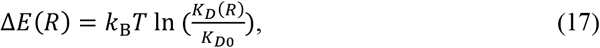

where 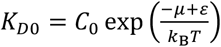. Because the numerical calculations cannot measure the complete binding energy Δ*G*(*R*) but only the membrane energy contribution, we calculate the relative energy change to compare with experiment:

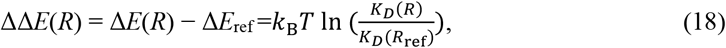

where Δ*E*_ref_ is chosen as the membrane energy change of the smallest vesicle (*R*_ref_ = 15 nm) in our experiments. With a constant excess concentration of unbound proteins and lipids across experiments as the curvature changes, this simplifies further to 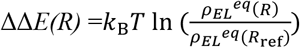.

### II.B Numerical implementation of the continuum membrane model

#### II.B.1. Finite element method for the continuum membrane model

In this work, we use three layers of triangular mesh to represent the bilayer membrane. The top and bottom layers of the mesh represent the neutral surface of each monolayer where the deformations of monolayer bending and area elasticity are energetically decoupled [50], and the middle layer represents the mid-surface of the bilayer membrane. The neutral surface is approximately located at the interface dividing the lipid polar heads and the hydrocarbon tails, and our leaflet height therefore quantifies the hydrocarbon length of each membrane leaflet, excluding the head groups [17].

To build the three layers of triangular mesh, we first build the outer monolayer mesh, and then move each vertex by the monolayer height *h*_0_ to get the mid-surface mesh, and move each vertex by 2*h*_0_ to get the inner monolayer mesh, with direction dependent on surface curvature. Each mesh layer thus has the same number of vertices and triangle elements, and the vertex indexes and triangle elements have a one-to-one correspondence. The distance between two triangle elements of the same index is projected onto the normal of the outer element to measure height.

To simulate the helix insertion on the outer monolayer surface, we select several adjacent triangles on the outer mesh to represent the location of the insertion, and the area sum of these selected triangles is the insertion size *a*. The spontaneous curvature of the selected triangles is set as *c*_0,*ins*_, the spontaneous curvature of the helix insertion, which differs from the lipid monolayer spontaneous curvature *c*_0_. The monolayer height has an equilibrium value 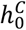, except at the insertion region, where the equilibrium height is decreased to 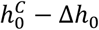. The equilibrium area of the outer monolayer is *A*_0,2_+ *a*, and the equilibrium area of the inner monolayer is *A*_0,1_. The values of 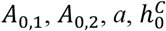 and Δ*h*_0_ remain constant during the energy minimization.

The total energy of the membrane is calculated by Eq (1) combined with terms Eq (2), Eq (3) and Eq (17), where we note the curvature energy of Eq (2) is defined by the integration on the equilibrium area of each monolayer [41]. In the finite-element analysis, the total energy for the bilayer membrane is defined as,

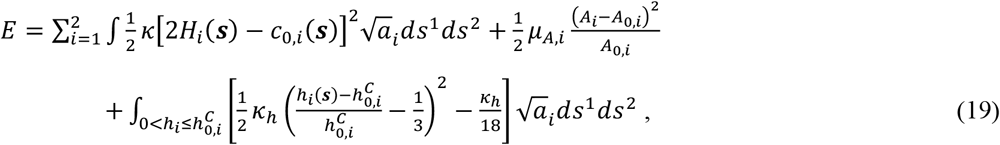

where the index *i* = 1, 2 refers to the inner and outer monolayer, respectively. {*s*^1^, *s*^2^} is the curvilinear coordinates on the curved surface, and ***s*** = ***s***(*s*^1^, *s*^2^) represents each point on the surface in three-dimensional space and the metric 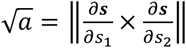. The spontaneous curvature varies along the surface, *c*_0_(***s***), following helix insertion. We integrate over the membrane surface numerically using second-order Gauss quadrature, which we showed is accurate [15]. To ensure robust numerical integration, we apply a regularization energy that keeps triangles within the mesh from significantly reshaping [15]. This regularization energy is not physical but purely technical to improve the numerical integration, and it converges to zero as the equilibrium state is reached. The force on each vertex on each of the three mesh layers is calculated as the gradient of the total energy due to curvature, area, tilt/height, and volume, see SI for detailed definitions. We increase the resolution of the mesh around the helix insertion to better capture the localized deformation driven by the insertion (see SI Methods), at a distance of <5nm showing good convergence of the energy (Fig S1). This increases the types of irregular patches on the membrane, which we account for in our numerical approach (Table S1). The equilibrated membrane energy and structure following helix insertion is calculated by performing energy minimization via gradient descent using both nonlinear conjugate gradients and steepest descent algorithms. For all parameter values, see Table 1. Unless otherwise noted, we use parameter value of the DOPC membrane as a default.

**Table 1.** Parameters of the continuum membrane model and values for 4 types of lipid membranes.

| Parameter | Symbol (unit) | DOPC | DLPC | POPC |
| --- | --- | --- | --- | --- |
| Bilayer hydrocarbon height | $2h_0$ (nm) | 2.71 [17] | 2.09 [19] | 2.75 [22] |
| Monolayer spontaneous curvature | $c_0$ ($\text{nm}^{-1}$) | -0.04 [54] | +0.11 [54] | +0.01 [54] |
| Bilayer bending modulus | $2\kappa$ ($\text{k}_\text{B}\text{T}$) | 19.4 [20] | 20.4 [20] | 25.7 [20] |
| Tilt modulus | $\kappa_\theta$ (pN/nm) | 89 [20] | 55 [20] | 73 [20] |
| Bilayer area modulus | $2\mu_A$ (pN/nm) | 265 [18] | 234 [55] | 255 [55] |
| Monolayer height modulus | $\kappa_h$ (pN/nm) * | 267 | 165 | 219 |
| Insertion spontaneous curvature | $c_{0,\text{ins}}$ ($\text{nm}^{-1}$) | 0.3 [15] | 0.4 [15] | 0.3 [15] |
| Decrease of the equilibrium height | $\Delta h_0$ (nm) * | 0.17 | 0.17 | 0.17 |

\*Note: The height modulus is $\kappa_h = 3\kappa_\theta$ . The equilibrium height decrease $\Delta h_0$ is validated by our experiments.

#### II.B.2 Boundary conditions of the continuum membrane

Our continuum membrane model can adopt arbitrary shapes, just as in our single-layer model[15, 56]. The vesicle membrane has an enclosed structure, whereas the tubule and flat membranes have edges that require imposed boundary conditions (BCs). Our model supports the same three types of boundary conditions implemented in our single-layer model [15]: free, fixed and periodic boundaries (Fig S2). In this paper we use periodic boundaries for the flat or tubular membranes. We create three rows of temporary or ‘ghost’ vertices outside the edge of the mesh with positions determined by translation of the mesh vertices at the opposite edge (Fig S2). The code is available open source at https://github.com/Yibenfu/triple-layer-continuum-membrane-model.

### II.C Molecular dynamics simulations

Each of the three systems (DOPC, POPC, DLPC) were built using the CHARMM-GUI web interface [57] with one helix peptide per leaflet and identical leaflet lipid counts (80 lipids per leaflet) such that the tension of the leaflets were equal (both at zero applied tension). Helix peptides were offset by half the box length in each lateral direction such that they were unlikely to interfere across leaflets. A one femtosecond timestep was used for dynamics. Three replicas of each bilayer without peptide were also modeled. Each system was run with the NAMD 2 simulation package [58] for 100 nanoseconds, which was sufficient to relax both the depth of the helix as well as the surrounding acyl chains, which relax on sub-nanosecond timescales[59]. Standard simulation parameters were applied consistent with the development of the forcefield (CHARMM C36m protein [60] and C36 lipid [61]): Lennard-Jones interactions were cutoff at 12 Angstroms, with force-switching applied starting at 10 Angstroms; the SHAKE/SETTLE algorithm constrained hydrogen bonds. The particle mesh Ewald algorithm computed long-range electrostatics with interpretation order six. Temperature was kept at 298K using a Langevin thermostat (1/ps damping coefficient). Isotropic pressure of 1 bar and zero lateral tension were maintained with a Langevin barostat (50 femtosecond oscillation period with 25 femtosecond decay time).

The shape of the surrounding membrane was computed using an algorithm initially described in Ref [62] to quantify lipid distortions around the (roughly angularly symmetric) transmembrane peptide gramicidin A, but adapted here to record leaflet deformations along a single axis. Briefly, lipids are sorted by their center-of-geometry position along an in-plane axis perpendicular to the long helix axis, with separate lists in the negative and positive directions (coordinate *d*). Only lipids within the end caps of the helix (+/-5 Angstroms), as measured by a second lateral coordinate perpendicular to the 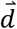 axis, were included. For each bin along this axis (0.6 Angstrom width), the average *d* and *z* value of each atom is stored, along with the number of lipids observed in the bin. Finally, starting from the peptide center, bins are combined to represent two total lipids. The set of *d, z* pairs for each averaged set of bins then represents the average trace (or shape) of two lipids in that region. Critically, the procedure assures that this trace always incorporates entire lipids into its average, even though it might average many lipids with wildly different shapes.

### II.D. In vitro experiments of ENTH on vesicles of increasing size

#### II.D.1. Experimental materials

DOPC (1,2-dioleoyl-*sn*-glycero-3-phosphocholine), POPC (1-palmitoyl-2-oleoyl-glycero-3-phosphocholine), DLPC (1,2-dilauroyl-sn-glycero-3-phosphocholine), and DOPI(4,5)P2 (1,2-dioleoyl-sn-glycero-3-phospho-[1’-myo-inositol-4’,5’-bisphosphate] [ammonium salt]) were purchased from Avanti Polar Lipids. DPPI(4,5)P2 (Phosphatidylinositol 4,5-bisphosphate diC16) was purchased from Echelon Biosciences. DP-EG10-biotin (dipalmitoyl-decaethylene-glycol-biotin) was generously provided by Darryl Sasaki from Sandia National Laboratories, Livermore, CA. Texas Red-DHPE (Texas Red 1,2-dihexadecanoyl-*sn*-glycero-3-phosphoethanolamine triethylammonium salt), NeutrAvidin, and Zeba spin desalting columns (7K MWCO, 5 mL) were purchased from Thermo Fisher Scientific. TCEP (tris(2-carboxyethyl) phosphine hydrochloride), PMSF (phenylmethanesulfonyl fluoride), EDTA-free protease inhibitor tablets, imidazole, PLL (poly-L-lysine), and ATTO-488 NHS-ester were purchased from Sigma-Aldrich. Sodium chloride, HEPES (4-2(2-hydroxymethyl)-1-piparazineethanesulfonic acid), IPTG (isopropyl-β-D-thiogalactopyranoside) and β-mercaptoethanol were purchased from Fisher Scientific. Amine reactive PEG (mPEG-succinimidyl valerate MW 5000) and PEG-biotin (biotin-PEG SVA MW 5000) were purchased from Laysan Bio, Inc. Glutathione Sepharose 4B was purchased from GE Healthcare. All reagents were used without additional purification.

#### II.D.2. Protein Purification and Labeling

ENTH was purified and labeled as previously described [1]. Briefly, ENTH was expressed as a fusion protein containing N-terminal glutathione-S-transferase (GST) in *E coli* BL21. These cells were lysed and centrifuged, and the protein was purified from the supernatant with affinity columns in a three-step process. First, the supernatant was incubated with glutathione agarose beads, and excess cell material was washed from the bound proteins. Thrombin was used to cleave the proteins from the beads, and excess thrombin was removed with *p-*aminobenzamidine-agarose. The purity of the protein was verified using SDS-PAGE and UV-Vis spectroscopy. The product was then concentrated and stored at −80C after snap freezing in liquid nitrogen. The purified protein was labeled with ATTO488 NHS-ester that was pre-dissolved in DMSO. The total DMSO in the protein/dye mixture was kept to less than 1.5% by volume. The mixture was incubated at room temperature for 30 minutes at a 2x stoichiometric excess of ATTO488 NHS-ester. Excess dye and DMSO were removed by running the reaction solution through a 7k Zeba Spin column.

#### II.D.3. Small unilamellar vesicle (SUV) Prep

Lipid aliquots were brought to room temperature and combined appropriately to achieve the following mixtures:

- 87.5% DOPC, 10% DOPI(4,5)P2, 2% DP-EG10-biotin, 0.5% Texas Red-DHPE
- 87.5% POPC, 10% DOPI(4,5)P2, 2% DP-EG10-biotin, 0.5% Texas Red-DHPE
- 87.5% DLPC, 10% DOPI(4,5)P2, 2% DP-EG10-biotin, 0.5% Texas Red-DHPE

All solvents for combined lipids were adjusted accordingly to yield a solution that was 65:35:4 Chloroform:Methanol:Water by volume. These solutions were evaporated under a nitrogen stream to create a lipid film and dried overnight under vacuum. The lipid film was rehydrated to a final lipid concentration of 100 uM in 25 mM HEPES (pH 7.4), 150 mM NaCl, 0.5 mM EDTA, and 0.5 mM EGTA. This lipid suspension was cycled three times through a freeze/thaw cycle (2 minutes to freeze in liquid nitrogen, and 2 minutes to thaw in a 60° C water bath). It was followed by extrusion through a 50nm and 200nm polycarbonate filter with average diameters of 79 nm and 127 nm, respectively, determined by dynamic light scattering. The two populations of SUVs were then mixed together to generate a broad range of vesicle diameters.

#### II.D.4. SUV Tethering and imaging

SUV’s were tethered as previously described [10]. Briefly, glass cover slips were cleaned and then passivated with a layer of biotinylated PLL-PEG. Vesicles with 2 mol% DP-EG10-Biotin were tethered to this surface using NeutrAvidin. Excess vesicles and NeutrAvidin were rinsed away. Labeled ENTH was then incubated with the tethered vesicles at the desired concentrations and imaged after a minimum of 10 minutes. Proteins and vesicles were visualized on a laser scanning confocal microscope (Leica Stellaris 5) using 488 and 561 nm lasers for excitation for the proteins and lipids, respectively.

## III. RESULTS

### III.A. Helix insertion introduces a local thinning of the membrane monolayer in all-atom MD simulations

We performed all-atom MD simulations of a membrane bilayer with a single ENTH helix inserted into the outer leaflet. The results show that the lipids tilt to encircle the sides and bottom of the helix (Fig 2). By characterizing the lipid heights as a function of distance from the insertion, we measured a decrease in the membrane height of approximately 1.12 Å for the DLPC membrane, with the perturbation extending about 2 nm from the insertion. The inner leaflet of the membrane arches slightly toward the insertion, as quantified in Fig 2. Our MD simulations with DOPC and POPC membranes yielded similar results, showing that the outer leaflet sinks around the insertion while the inner leaflet arches toward it. However, the decrease in leaflet height varies with lipid composition. For the DOPC membrane, the observed decrease in membrane height is about 0.66 Å (Fig S3A), while for the POPC membrane, the observed height decrease is approximately 0.86 Å (Fig S3B). Despite the larger deformation of the DLPC membrane following helix insertion, we show from in vitro experiment and continuum modeling below that it shows reduced curvature sensing based on multiple changes to its material properties.

**Figure 2.**
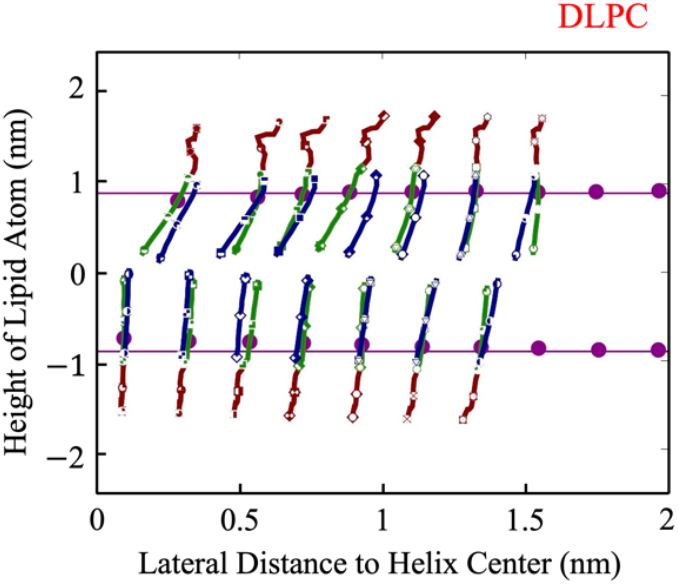
MD simulations captured the lipid trace around the ENTH helix, located at coordinates (0,1), with the DLPC membrane. The sn-1 chain is depicted in blue, and the sn-2 chain in green. The extended headgroup region is represented in dark red. The CHARMM C24 atom, marking the approximate neutral surface, is shown by the purple dots, with the height of the unperturbed membrane shown by the purple line. DOPC, POPC membranes in Fig S4.

### III.B. Bilayer continuum membrane model captures local thinning around the insertion

We find that enforcing a change to the equilibrium height at the insertion patch (Δ*h*_0_) to mimic the lipid tilt observed in the MD simulations drives a local thinning of the leaflet in the continuum model, in agreement with the MD simulations (Fig 3). The thinning is driven both by the height reduction in the outer leaflet around the helix insertion, as well as a compensatory height loss in the inner leaflet due to the coupling between the leaflets, also in agreement with the MD simulations. Both perturbations dampen with radial distance from the insertion as the membrane returns to its equilibrium height within 2∼4 nm of the insertion, Fig 3A. We find that both the helix spontaneous curvature *c*_0,*ins*_ and the value of Δ*h*_0_ contribute to the local membrane deformation of both leaflets around the insertion. As Δ*h*_0_ increases with fixed *c*_0,*ins*_, the outer leaflet sinks deeper near the insertion and both the inner leaflet and the middle surface arch higher, resulting in increased thinning of the membrane (Fig 3B). The sinking of the outer leaflet around the insertion is not monotonic with distance, as it is accompanied by an arch upward before relaxing to the equilibrium height (Fig 3B). In contrast, increasing *c*_0,*ins*_ while keeping Δ*h*_0_ fixed systematically shifts up the displacement of all leaflets, such that the overall thinning of the membrane is of similar magnitude (Fig 3C). Therefore, optimizing agreement between this structural deformation produced by MD data and the continuum model requires selection of coupled Δ*h*_0_ and *c*_0,*ins*_ values, of which more than one pair shows good agreement, and no pair is in perfect agreement. The best fit values of both are chosen to minimize the error between the MD results and the in vitro experimental results, as described below. For DLPC we use Δ*h*_0_ = 0.17*nm* and c_0,*ins*_ =0.4 nm^−1^ (Fig S3 and Table 1). We note the membrane thickness *h*_0_ can also influence the leaflet deformation (Fig S4), but this parameter is fixed to its known value for each lipid (Table 1).

**Figure 3.**
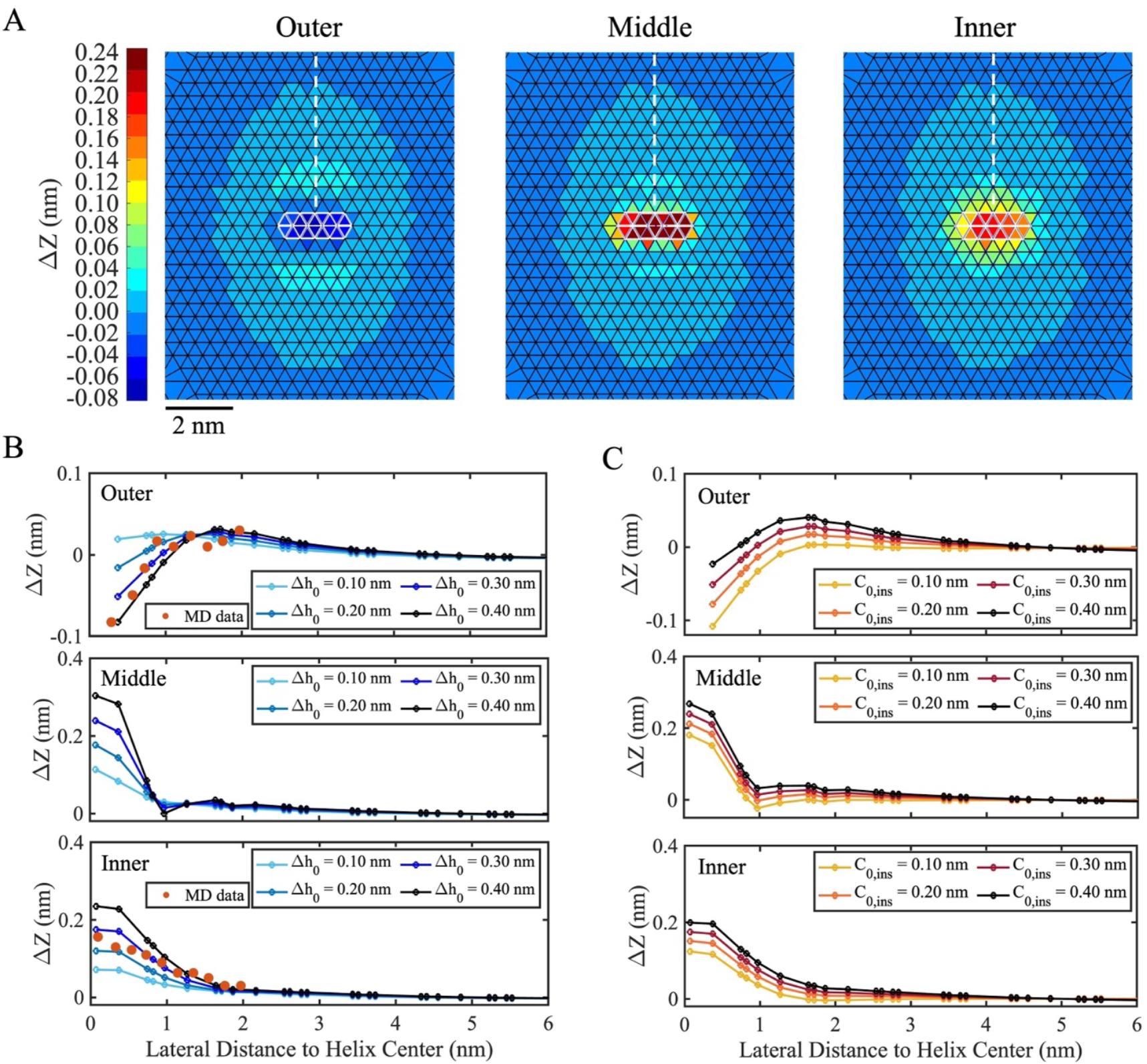
The DLPC lipid membrane is deformed due to the helix insertion, demonstrated by our continuum membrane model. The membrane is initially flat and positioned in the *X*-*Y* plane. The height is represented by the *Z* coordinate of each triangle element center, with Δ*Z* defined as Δ*Z* = *Z* – *Z*_0_, where *Z*_0_ is the height far from the helix center. (A) The radial height deformation of the membrane around the helix insertion impacts the outer leaflet (left panel), middle surface (middle panel), and inner leaflet (right panel). The white patch indicates the location of the helix insertion on the outer leaflet, and the corresponding regions on the middle surface and inner leaflet, with a more extended perturbation observable on the axis perpendicular to the helix axis. (B) The height deformation of the membrane decreases with increasing distance from the helix insertion, depending on Δ*h*_0_, where *c*_0,*ins*_ = 0.3 nm^−1^ is fixed. The membrane height is calculated on triangle centers located along the while dashed line shown in (A). The red dots represent MD data from Fig 2. To compare the continuum membrane model results with the MD data, the red dots are shifted vertically to best align with the continuum membrane model’s height. Note that the lateral position of the MD data on the outer leaflet does not start until ∼0.2nm due to the volume occupied by the helix, whereas inner leaflet extend to 0 nm. (C) The shape of the membrane is perturbed by the helix insertion dependent on *c*_0,*ins*_, with Δ*h*_0_= 0.3 nm fixed.

### III.C. The bilayer continuum model captures curvature sensing

Consistent with the single-layer continuum membrane model, we here validate that the bilayer continuum model also reproduces curvature sensing by amphipathic helices. Specifically, we performed simulations with one *α*-helix inserted on a vesicle across a range of vesicles with increasing radii and thus decreasing curvature. We find that the membrane energy change Δ*E* due to one helix insertion increases monotonically as the vesicle becomes larger and starts to plateau as the curvature increases towards a flat membrane, as observed in multiple experiments, indicating the most stable interaction for the most highly curved membranes (Fig 4A). By analyzing the three components of the energy-- the curvature energy, the area elasticity energy, and the height elasticity energy--we found they all increase for larger vesicles, indicating that all three contribute positively to the curvature sensing phenomenon. By re-zeroing each energy difference to the value at the smallest vesicle *R* = 15, ΔΔ*E* shows that the curvature energy and area elasticity energy increase faster than the height elasticity energy with increasing vesicle size (Fig 4B). Thus these terms are more dominant in controlling the curvature sensitivity of the *α*-helix insertion.

**Figure 4.**
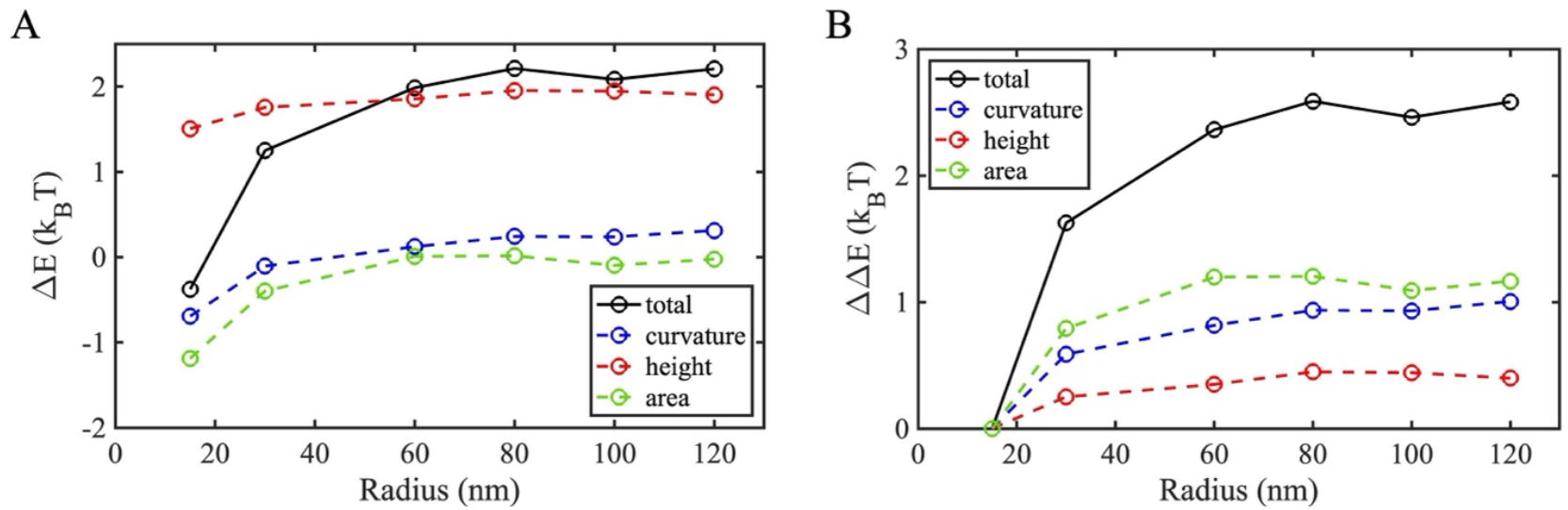
Bilayer continuum model captures curvature sensing by α-helix insertion. (A) Membrane energy change relative to the unperturbed vesicle following one α-helix insertion is measured vs the vesicle radius. Black is total energy change, with individual contributions plotted from curvature (blue), area (green), and height (red). (B) By re-zeroing the membrane energy of (A) by the reference value at R=15nm we plot ΔΔ*E*(*R*) *=* Δ*E*(*R*)−Δ*E*(R=15nm). Simulations were performed with DOPC membrane parameters with the bilayer hydrophobic height *h*_0_ = 5.0 nm, helix insertion *a* =2 nm^2^, *c*_0,*ins*_ = 0.3 nm^−1^ and Δ*h*_0_ = 0.17 nm.

### III.D. Model reproduces curvature sensing measured *in vitro* for varying lipid compositions

To test our model predictions, we performed *in vitro* experiments measuring curvature sensing of the ENTH domain to vesicles of varying radii, as in previous work [10], but here with variations to lipid composition. To eliminate changes to chemistry that could influence recruitment of ENTH to the membranes, we kept the PIP_2_ concentration the same and compared DOPC, POPC and DLPC containing liposomes. Thus, the head group phosphatidyl choline remained unchanged but the tail groups varied. The expected material parameters for these lipids (e.g. bending and tilt moduli) are known and collected in Table 1. Because the tail groups influence all material parameters, it is not obvious how the curvature sensing of ENTH to highly curved membranes will vary. From our experiments, we see that ENTH shows stronger curvature sensing, or a higher relative affinity for highly curved membranes, when they are composed of DOPC compared to DLPC (Fig 5). The change in binding energy between the largest and smallest vesicle for DOPC reaches nearly 2.0 k_B_T, while for DLPC membrane it is closer to 1.5 k_B_T. POPC is similar to DOPC but with slightly stronger curvature sensing.

**Figure 5.**
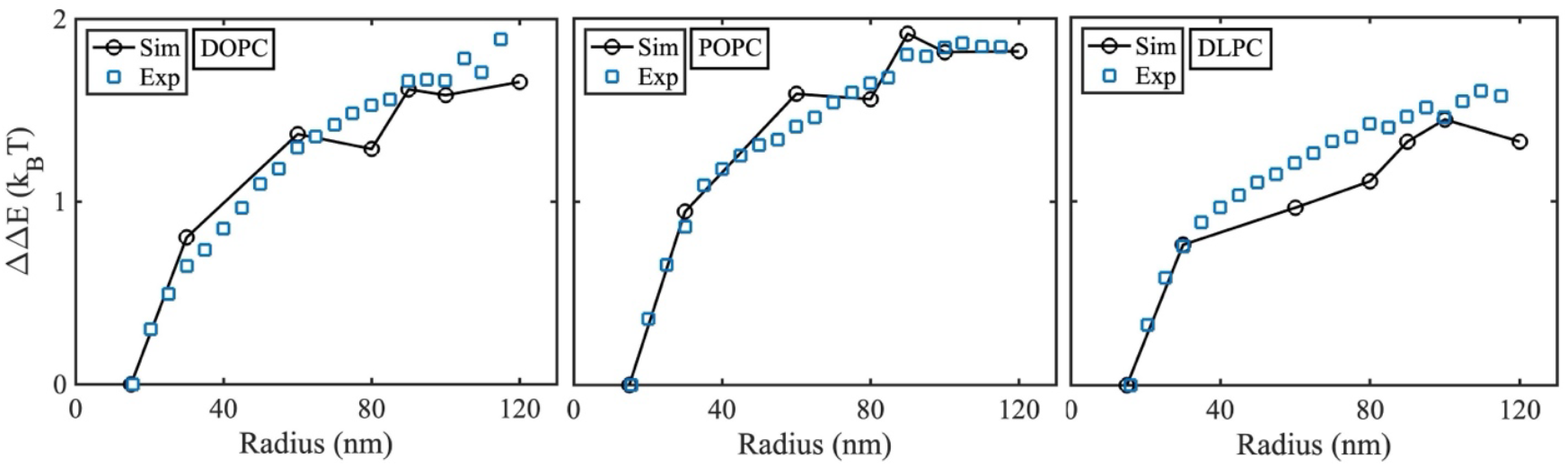
Bilayer continuum model can reproduce and explain curvature sensing observed in in vitro experiments. (A) ENTH helix binding to membrane of DOPC lipid. (B) ENTH helix binding to membrane of POPC lipid. (C) ENTH helix binding to membrane of DLPC lipid. The parameters in the simulations are as in the Table 1. The experiments were performed with the ENTH concentration 100 nM.

We mimicked these same symmetric lipid compositions with our bilayer continuum model using parameters in Table 1 and the same vesicle radii as in the experiment, finding strong agreement with experiment (Fig 5). For the helix insertion, the only remaining parameters not specified from the literature are *c*_0,*ins*_ and Δ*h*_0_, which must be constrained based on the observed deformation from the MD simulations. However, more than one pair of *c*_0,*ins*_, Δ*h*_0_ values show agreement with deformations in the upper and lower leaflet (Fig 3, Fig S3). We found that using the same values of *c*_0,*ins*_ for DOPC and POPC membranes, but a slightly larger value of *c*_0,*ins*_ for the thinner DLPC is effective and consistent with previous work showing higher *c*_0,*ins*_ on thinner membranes[31]. We tested variations in the insertion depth Δ*h*_0_ for each lipid type within a range of values consistent with our MD simulations (Fig 2, Fig S3), but we found good agreement for all membranes using Δ*h*_0_ = 0.17 nm.

From our continuum model, we can explain the reduced degree of curvature sensing when switching from DOPC to DLPC, despite having nearly identical bending modulus. As we systematically characterize below, a more positive spontaneous curvature of the lipid *c*_0_ (from −0.04 to 0.11) and a thinner height of the lipid (from 2.71 to 2.09) when going from DOPC to DLPC will both reduce curvature sensing. Although DLPC has a larger deformation following helix insertion (Fig 2, Fig S3), likely due to its lower tilt modulus (Table 1), this property is offset by the changes to its height and shape (*c*_0_) compared to DOPC and POPC. A slight increase in POPC curvature sensing compared to DOPC is expected given its higher bending modulus [15], which outweighs the more modest increase from *c*_0_=-0.04 to 0.01.

### III.E. Thicker membrane enhances membrane binding and curvature sensing

Our results show that by increasing the membrane height while keeping the vesicle size fixed, Δ*E* due to insertion decreases (Fig 6A), indicating that the helix insertion is more stable on thicker membranes. By the comparison of the three energy components, we see, as expected, that the height elasticity energy contributes the most to sensing of the membrane height, although the curvature energy also decreases with thicker membranes. The same trend is conserved for small and large vesicles (Fig 6B). Finally, by again comparing the relative energy change vs the R=15nm vesicle, we see that the relative increase or sensing of the curvature is highest for the thicker membranes (Fig 6C and D). The insertion is relatively more stable for small and thicker vesicles, and relatively less stable for large and thinner membranes.

**Figure 6.**
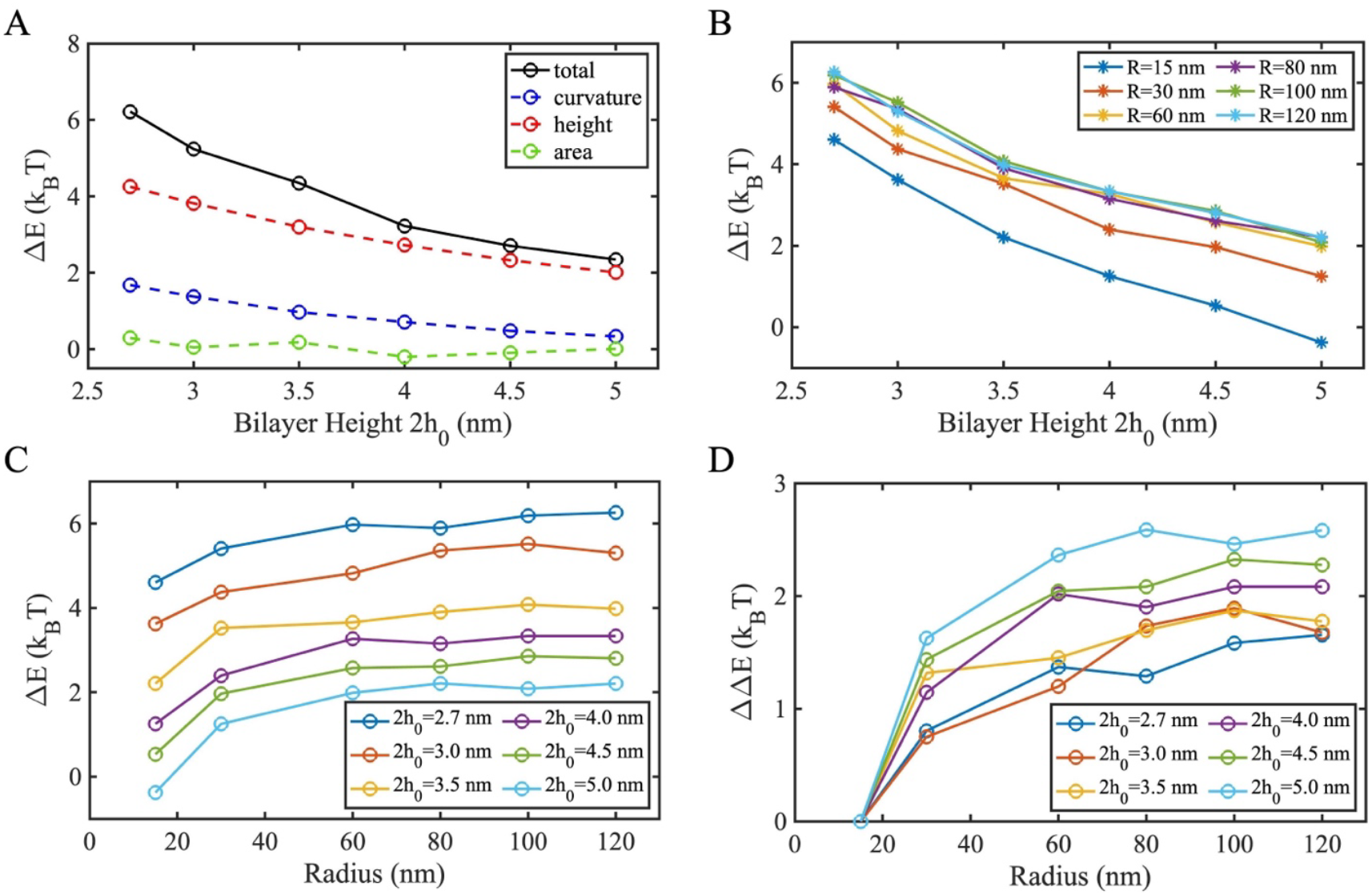
Thicker membranes produce increased curvature sensing. (A) Membrane energy change Δ*E* because of one α-helix insertion vs membrane height 2*h*_0_ and contributions from curvature (blue), height(red) and area (green). Simulations were performed with vesicle radius *R* = 90 nm. (B) Membrane energy change Δ*E* vs 2*h*_0_ on vesicles of different radii. (C) Membrane energy change Δ*E* vs vesicle radius shows curvature sensing at all different heights (2*h*_0_) tested. (D) Re-zeroed energies from (C) (ΔΔ*E(*R*)* = Δ*E*(*R*)−Δ*E*(*R*=15nm). Simulations were performed with DOPC membrane parameters, *a* =2 nm^2^, *c*_0,*ins*_ =0.3 nm^−1^ and Δ*h*_0_ = 0.17 nm.

### III.F. Deeper helix insertion strengthens curvature sensing

Distinct *α*-helices can insert into the membrane at varying depths depending on their sequence and the membrane composition[63]. With a deeper insertion of the helix into the membrane, the lipids will have to deform and tilt more significantly around the insertion, which in our model is captured by a larger Δ*h*_0._ We find that larger Δ*h*_0_ increases Δ*E*, meaning that the mechanical energy is destabilized more by a deeper insertion (Fig 7A). Similar to the dependence on membrane thickness, the energy sensitivity due to changes in Δ*h*_0_ is predominantly due to the height elasticity, with smaller contributions due to curvature. Deeper insertions lead to stronger curvature sensing (Fig 7B), as the difference in Δ*E* between binding to small and large vesicles is larger. We note that even if Δ*h*_0_=0, the bilayer model still senses curvature but to a lesser degree. This is because of the significant contribution of curvature and area elasticity energy towards curvature sensing. However, setting Δ*h*_0_=0 does not reproduce the structural deformation of the membrane as demonstrated by the MD simulations (Fig 2, Fig S4).

**Figure 7.**
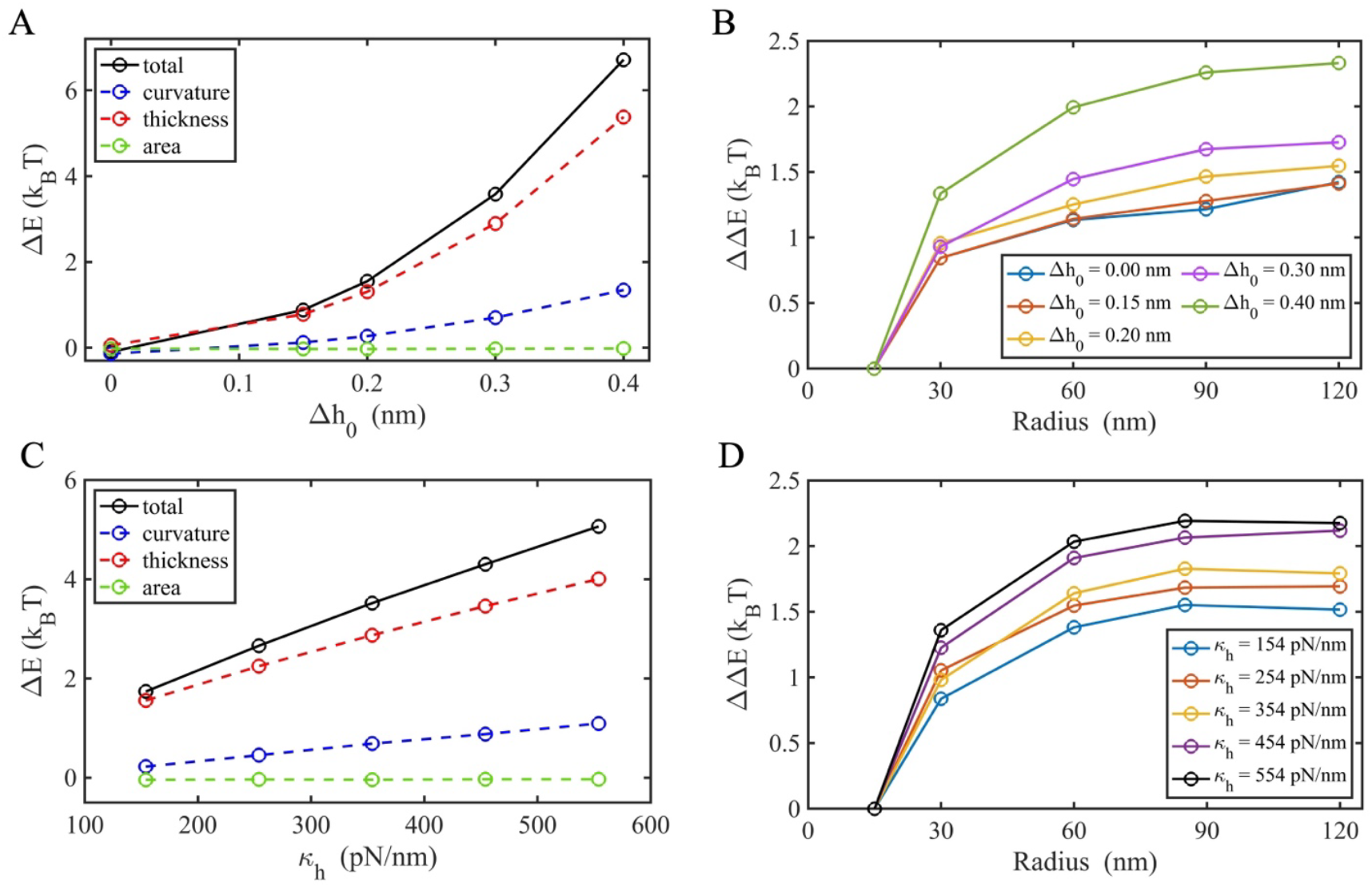
Helix insertion depth and lipid tilt modulus affects curvature sensing. (A) Membrane energy change Δ*E* because of one α-helix insertion depends on the equilibrium value of the height decrease at the insertion Δ*h*_0_. Simulations were performed with vesicle radius *R* = 60 nm. (B) Relation of ΔΔ*E* and vesicle *R* is regulated by Δ*h*_0_. Here, ΔΔ*E=* Δ*E*− Δ*E*(R=15nm).

### III.G. The spontaneous curvature of the membrane leaflets modulates curvature sensing in symmetric bilayers

Each leaflet of a membrane bilayer will have a spontaneous curvature *c*_0_ that is determined by its lipid composition, mostly because of the lipid’s shape but also due to interactions between neighboring lipids[25] (e.g. DOPC has *c*_0_ =−0.04 nm^−1^, and DLPC has *c*_0_ =+0.11 nm^−1^; Table 1). If the two leaflets are symmetric in their lipid composition, this results in a bilayer spontaneous curvature of zero, meaning the bilayer prefers to be flat. However, the helix only inserts into one of the leaflets, and previous experimental work on symmetric bilayers showed that lipid compositions with a more negative *c*_0_ induce stronger binding of the helix-inserting domain of Arf1[13, 32]. Our model reproduces this same result here, with very good agreement to the experiment (Fig 8), showing that as we decrease *c*_0_ from −0.06 nm^−1^ to −0.25 nm^−1^, Δ*E* decreases primarily due to the contribution from the curvature energy, indicating a more stable membrane energy (Fig 8A). This result reinforces that the mechanical energy of the membrane that varies with lipid composition helps determine how strongly an inserted helix will bind to the membrane.

**Figure 8.**
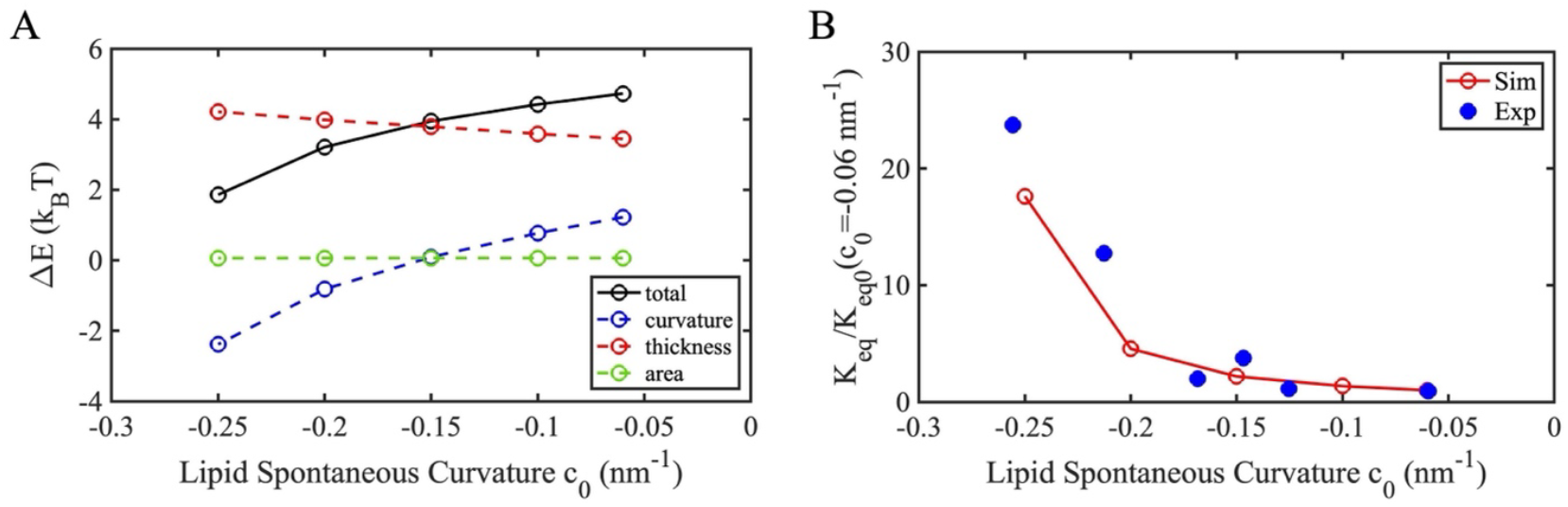
Helix insertion senses the lipid spontaneous curvature *c*_0_. (A) Membrane energy change Δ*E* because of one α-helix insertion depends on the lipid spontaneous curvature *c*_0_, which effectively reports on the preferred shape of the lipid (cylinder, wedge, or inverted wedge) (B) The calculated relation of *K*_*eq*_/*K*_*eq0*_ with *c*_0_. Here, simulations were performed with the flat membrane, bilayer height 2*h*_0_ = 2.71 nm, bilayer area modulus 2*μ*_*A*_ = 250 pN/*μ*s, bilayer bending modulus 2**κ** = 25 k_B_T, height modulus **κ**_*h*_*=*267 pN/nm, insertion spontaneous curvature *c*_0,*ins*_ =0.3 nm^−1^ and Δ*h*_0_ =0.2 nm.The helix insertion size is 4 nm^2^ (4 nm in length and 1 nm in width), consistent with the Arf1 helix size. The experiment data for the helix-inserting domain of Arf1 is sourced from Ref [13, 32]. We converted our Δ*E* to a relative binding equilibrium constant using *K*_*eq*_/*K*_*eq0*_ = exp[− (Δ*E* − Δ*E*_0_)/*k*_B_*T*], where Δ*E*_0_ is the membrane energy change on the reference membrane with the spontaneous curvature *c*_0_ = −0.06 nm^−1^, and *K*_*eq0*_ is thus the corresponding equilibrium constant for this reference membrane. Both simulation and experiment show a sharp increase in *K*_*eq*_/*K*_*eq0*_ as *c*_0_ becomes more negative.

### III.H. Helix insertion does not sense membrane asymmetry

The plasma membrane is actively maintained with a significant asymmetry in membrane composition across the inner and outer leaflet [36]. While generating membrane asymmetry is thus critical for membrane function in living systems, we specifically query here whether the mechanical consequences of asymmetry, as parameterized by distinct leaflet spontaneous curvatures, would impact curvature sensing by helix insertion. With one helix inserted on the outer leaflet of the flat membrane, we varied the spontaneous curvature of the inner leaflet *c*_0,1_ while keeping the outer leaflet value *c*_0,2_constant at zero. We find that the membrane energy change is quite marginal even as *c*_0,1_ varies from −0.2 to 0.2 nm^−1^ (Fig 9A). This result holds even as we change the insertion depth of the helix Δ*h*_0_, showing some fluctuations in energy values (<0.5 k_B_T) that do not report on any consistent trend with either Δ*h*_0_ or *c*_0,1_ (Fig 9B). Therefore, we conclude that the helix insertion is not strongly coupled to the spontaneous curvature of the opposing leaflet, but only to the spontaneous curvature of its inserted leaflet, as quantified in the previous section (Fig 8).

**Figure 9.**
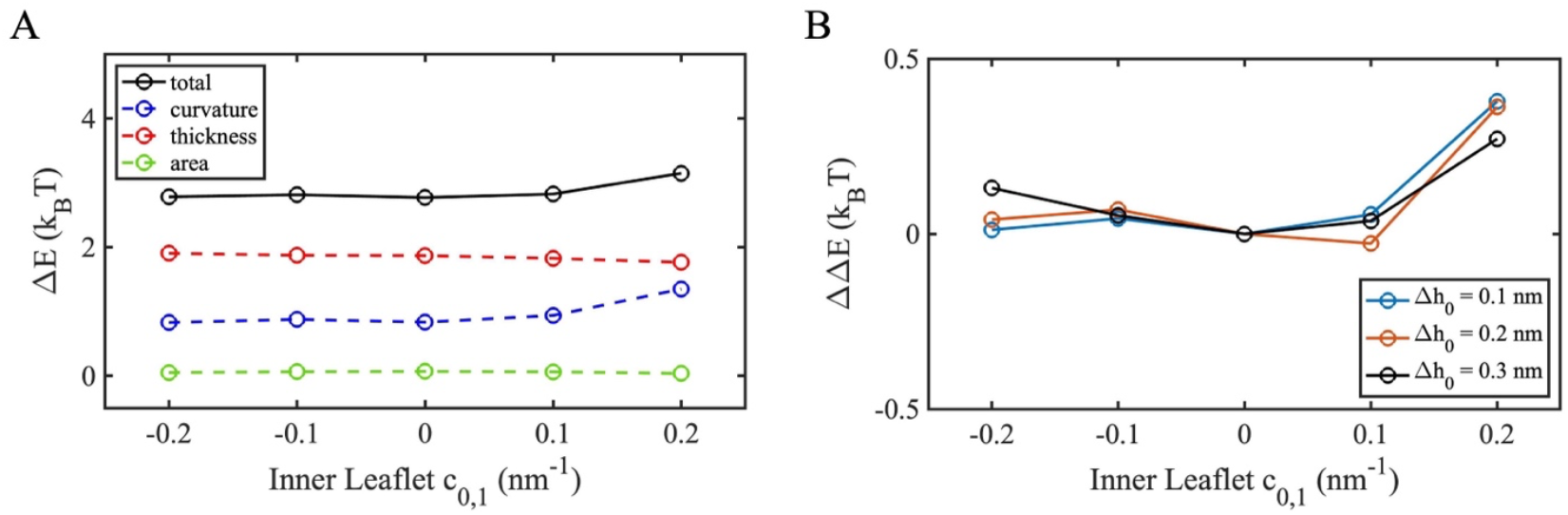
Helix insertion does not sense membrane asymmetry. (A) The membrane energy change Δ*E* following helix insertion is not sensitive to the lipid spontaneous curvature *c*_0,1_ of the inner monolayer, while the outer monolayer has a fixed *c*_0,2_ =0. (B) Changes to insertion depth Δ*h*_0_ also do not systematically impact the energetics of helix insertion as a function of *c*_0,1._ Here, ΔΔ*E=* Δ*E*− Δ*E*(*c*_0,1_ = 0 nm^−1^). The simulations were performed with a flat membrane, the helix insertion on the outer monolayer is 2 nm^2^ (2 nm in length and 1 nm in width). The membrane parameters follow those of the DOPC membrane, except for *c*_0_ and Δ*h*_0_.

## IV. Discussion

Our results demonstrate how continuum membrane models that capture key mechanical properties of the membrane including bending modulus, tilt modulus, area compressibility, and bilayer height can accurately capture both structural and energetic changes to the bilayer upon amphipathic helix insertion, as measured here by direct comparison to all-atom MD simulations and in vitro experiments on the ENTH helix. Our model recapitulates the stronger curvature sensing measured here experimentally for membranes that are identical except for the tail groups of the PC head-group lipids changing from DLPC to DOPC and POPC. Notably, DLPC and DOPC have the same bending modulus, so we know that is not changing the curvature sensing[15]. We learned from our model here that the primary driving force for this difference is the height and shape change of DLPC as parameterized by *h*_0_ and *c*_0_, both of which select for reduced curvature sensing (Fig 6, Fig 8). The larger deformation in DLPC observed in the MD simulations on flat membranes (Fig 2, Fig S3) is not sufficient to energetically counterbalance the changes to *h*_0_ and *c*_0_. Predicting these responses *a priori* would not be possible due to the competing and nonlinear dependence of the membrane mechanical energy and shape changes on these material parameters. We systematically assessed how variations in each material parameter of an explicit double-leaflet bilayer will change the curvature sensing by proteins inserted into the outer leaflet, which is not physically accessible experimentally or tractable using MD simulations. Similar to the single-surface continuum model [15], we again find that the mechanical energy due to helix insertion produces Δ*E* < 0 on the smallest vesicles (Fig 4), indicating that the insertion actually relieves stress in the highly curved small vesicles, as they are far from their preferred flat curvature. We showed how increasing the height of the bilayer or the negative spontaneous curvature of both leaflets also produce stronger curvature sensing (like DOPC), in excellent agreement with previous experimental work[13]. Somewhat surprisingly, asymmetry between the leaflets has negligible impact on curvature sensing, indicating the phenomenon is primarily driven by the properties of the outer leaflet where the helix inserts.

By incorporating both leaflets into our continuum model and thus expanding the material description of the bilayer, our results demonstrate that although lipid tilt/height contributes to the mechanical energy of the membrane following helix insertion, favoring more stable binding to more curved membranes, the largest contributors to curvature sensing are the bending and area energy. This is not to say that the tilt/height energy does not respond to the insertion, as particularly for DLPC we see an increase in the local membrane deformation following insertion, but that the energy change is not as markedly dependent on this local membrane curvature. This implies that even a single-surface continuum model provides a strong estimate of curvature sensing despite lacking explicit height variables[15]. Similarly, our results indicate that when it comes to curvature sensing, the coupling between the leaflets has a marginal impact even with significant variations in lipid shape compositions in distinct leaflets (Fig 9). Our model captures leaflet coupling via the enforcing of the incompressible membrane (e.g. 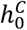 reflects the equilibrium height due to curvature-induced area changes to maintain a constant volume), and via the tilt/height term. This suggests that single-surface models can effectively model the properties of the outer leaflet when evaluating relative energy changes due to helix insertion. However, we expect that leaflet asymmetry would contribute much more to variations in membrane energy for an insertion or transmembrane protein that spanned both leaflets [43]. Thus another useful application of our model is in quantifying the energetic benefit of the shape preference and spatial localization of transmembrane proteins (alone or in clusters) within asymmetric membranes such as the plasma membrane[36]. Transmembrane proteins show a clear height preference to match their hydrophobic interiors to the hydrocarbon tail region of bilayers [43], with hybrid continuum-atomistic simulations demonstrating how the membrane deforms to stabilize interactions with ion channels [64, 65] or drive partitioning of lipids based on bilayer thickness[66]. Leaflet asymmetry is now achievable within *in vitro* experiments [36] and our model can efficiently predict how asymmetry in both leaflet height and spontaneous curvature determine the mechanical membrane energy around proteins.

Our bilayer continuum model is significantly more efficient than all-atom MD at measuring large-scale membrane shape (e.g. *μ*m) and energy changes even due to perturbations at the nanometer scale, with energies at the scale of a few k_B_T. For the large vesicles (R=150nm), the continuum calculations can optimize energy and shape in a few days to compare directly with experiment (Fig 5), whereas free energy calculations on vesicles of this size are not tractable in MD. Further, many of the membrane material properties needed for our continuum model (Table 1) can be found within the literature[20]. A significant limitation of the continuum model is it cannot predict directly from structure or sequence how the membrane will locally deform in response to insertions, and it lacks molecular-level details on lipid packing and reorganization around the helix insertion. Hence, we relied here on new MD simulations (Fig 2,3, Fig S3) to provide this input to the continuum model to help constrain parameter estimates for *c*_0,*ins*_, Δ*h*_0_. The continuum model also cannot predict a full binding free energy Δ*G*, as it computes only the mechanical contribution to the energetics. Another trade-off with any continuum approach, particularly compared to MD, is that it does not dynamically capture inhomogeneities in membrane material properties that would be driven by lipid diffusion, or flip-flop between leaflets (albeit over slow timescales) or possible phase separation of lipids within the bilayer[67]. While we can impose variations across our surface by representing material properties as scalar fields like *c*_0_(***s***), it best represents a steady-state compositional state of a membrane that will then respond mechanically and structurally to proteins or other external perturbations.

With insight here on the range of structural perturbations following helix insertion and the scale of the energy changes, we can anticipate how multiple helix insertions might induce cooperativity or feedback in recruitment to curved membranes. From the MD simulations and the continuum model, the helix perturbs the membrane only to ∼2 nm from the center of the helix. For ENTH, the helix is attached to a folded protein domain that will occupy a footprint of ∼4 nm by 1.2 nm. This suggests that even two adjacent ENTH proteins will have minimal overlap in the perturbation to the membrane. This supports the assertion that the helices are effectively independent of one another (see Methods), with each contributing the same Δ*E* to the membrane energy. This is further supported by our experimental data on curvature sensing by ENTH that indicates that as the total equilibrium concentration of proteins on the membrane increases, the measured equilibrium constant does not change systematically (Fig S5). In other words, there is no measurable evidence of cooperativity via membrane mechanics for the ENTH binding to vesicles, although a small effect would not be detected given the noise in experimental measurements. Previous work that explicitly measured membrane energies and deformations due to two adjacent insertions using classical elasticity theory did measure a lowering of the energy, or a cooperative effect, as the helices come close together [31, 68]. The energetic effect was quite small for peptides of finite (<6nm) length[68], while the deformations were significant in the full solid elastic model when the insertions on a cylinder were infinitely long[69]. Longer helices induce larger changes to the membrane energy and membrane structure, enhancing curvature sensing as shown by us and others [15, 35], which would therefore be expected to propagate perturbations to larger distances from the insertion center and be more likely to affect neighboring protein insertions. Together, our results here, along with these previous studies[15, 68, 69] predicts that long helices would be a necessary element for inducing a strong enough perturbation to generate experimentally measurable cooperativity in membrane-mediated binding. Such cooperativity would favor local short-range interactions and alignment between proteins, although we note this effect may be dwarfed by favorable protein-protein binding interactions between full-length helix-containing proteins like alpha-synuclein.

We provide our software open-source at github.com/yibenfu/triple-layer-continuum-membrane-model to facilitate extensions and applications of this model which accommodates diverse membrane geometries (including easily initialized red-blood cell shapes[15]), boundary conditions, and local or global changes to material properties as defined from published experiments (Table 1). Although this model predicts energetics and structure without dynamics, our continuum models can capture deformations driven by single proteins or rigid macromolecular assemblies[56]. Coupling this surface model to particle-based reaction diffusion on surfaces[70] and as coarse-grained proteins recruit to and assemble on membranes[9, 71, 72] will enable predictive timescales for membrane remodeling and the role of protein-protein interactions in overcoming kinetic barriers to membrane remodeling. As it is, this experimentally validated tool provides quantitative and testable predictions detailing energetic and structural changes to membranes as driven by proteins, as a key step in diverse membrane remodeling pathways throughout biology[23].

## Author Contributions

YF and MEJ designed the study. YF, AJS, and MEJ developed the continuum model. YF implemented the model and collected the continuum model results and figures. AHB and AJS performed and analyzed MD simulations. DHJ and WFZ performed all in vitro experiments. YF and MEJ wrote the initial manuscript. YF, DHJ, WFZ, AJS, and MEJ revised the manuscript. All authors approved the manuscript.

## Acknowledgements

M.E.J. gratefully acknowledges funding from a National Institutes of Health MIRA Award R35GM133644. We acknowledge use of the ARCH supercomputer rockfish at Johns Hopkins. AJS and AHB were supported by the intramural research program of the *Eunice Kennedy Shriver* National Institute Child Health and Human Development at the NIH. WFZ and DHJ gratefully acknowledge funding from a National Institutes of Health MIRA Award R35GM147333 to WFZ.

