## Supplemental Information for "Predicting protein curvature sensing across membrane compositions with a bilayer continuum model"

### Supporting Information
### for

#### Predicting how lipid composition controls protein curvature sensing with a bilayer continuum membrane model

Yiben Fu[1,2,3,4], David H. Johnson[5], Andrew Beaven[6], Alexander J. Sodt[6], Wade F. Zeno[5], Margaret E. Johnson[7]

*[1]School of Biomedical Sciences and Engineering, South China University of Technology, Guangzhou International Campus, Guangzhou 511442, P. R. China*

*[2]National Engineering Research Center for Tissue Restoration and Reconstruction, South China University of Technology, Guangzhou 510006, P. R. China*

*[3]Guangdong Provincial Key Laboratory of Biomedical Engineering, South China University of Technology, Guangzhou 510006, P. R. China*

*[4]Key Laboratory of Biomedical Materials and Engineering of the Ministry of Education, South China University of Technology, Guangzhou 510006, P. R. China*

*[5]Mork Family Department of Chemical Engineering and Materials Science, The University of Southern California, Los Angeles, California, 90089, USA*

*[6]Eunice Kennedy Shriver National Institute of Child Health and Human Development, National Institutes of Health, 9000 Rockville Pike, Bethesda, Maryland 20892, USA*

*[7]T. C. Jenkins Department of Biophysics, The Johns Hopkins University, 3400 N. Charles St., Baltimore, Maryland 21218, USA*

#### I. Derivation of the monolayer equilibrium area

From the main text, we defined the equilibrium areas of each monolayer $A_{0,i}$ relative to the mid-surface area $A_m$ by introducing a parameter $\gamma$. It can be solved for via $\frac{\partial E_{1+2}}{\partial \gamma} = 0$. The sphere and cylinder both have constant surface curvature $c_i$ and thus the formula for $E_{1+2}$ simplifies to

$$E_{1+2} = \frac{1}{2}\kappa(2c_1 - c_0)^2 A_{0,1} + \frac{1}{2}\mu_A \frac{\left(A_1 - A_{0,1}\right)^2}{A_{0,1}} + \frac{1}{2}\kappa(2c_2 + c_0)^2 A_{0,2} + \frac{1}{2}\mu_A \frac{\left(A_2 - A_{0,2}\right)^2}{A_{0,2}}. \quad (S1)$$

To derive $\gamma_{\text{sphere}}$, we consider a sphere of radius $R$, $A_m = 4\pi R^2$, $A_1 = 4\pi(R-h)^2$, and $A_2 = 4\pi(R+h)^2$. The mean curvatures of the leaflets are $c_1 = 1/(R-h)$ and $c_2 = 1/(R+h)$. By inserting these into Eq. S1 along with Eq. 4 from the main text and minimizing with respect to $\gamma$, we get:

$$\frac{(R-h)^4}{R^4\left(1-\gamma_{\text{sphere}}\right)^2} - \frac{(R+h)^4}{R^4\left(1+\gamma_{\text{sphere}}\right)^2} = \frac{\kappa}{\mu_A}\left[\left(\frac{2}{R-h} - c_0\right)^2 - \left(\frac{2}{R+h} + c_0\right)^2\right]. \quad (S2)$$

Eq. S2 shows that $\gamma_{\text{sphere}}$ is dependent on geometric and material parameters. The analytical expression is the root of a quartic equation [1]. For intuition, we simplify by setting the right-hand side to zero, which is satisfied if: (i) The monolayer has $c_0 = 2h/(R^2 - h^2)$; or (ii) $R$ is large, resulting in:

$$\gamma_{\text{sphere}} \approx \frac{2Rh}{R^2 + h^2} \quad (S3)$$

Now the monolayer areas are only dependent on the membrane geometry. As expected, with flatter membranes, $R \rightarrow +\infty$, and $\gamma_{\text{sphere}} \rightarrow 0$. The difference between the two monolayers' equilibrium areas in this limit is $\Delta A_0 = A_{0,2} - A_{0,1} = 16\pi R^3 h/(R^2 + h^2)$. For the large radius $R \gg h$, we have $\Delta A_0 = 16\pi R h$, which is the same as the prediction of the ADE model [2]. For a cylindrical membrane with length $L$, and radius $R$, $A_m = 2\pi R L$, $A_1 = 2\pi(R-h)L$ and $A_2 = 2\pi(R+h)L$. The mean curvatures are $c_1 = 1/2(R-h)$ and $c_2 = 1/2(R+h)$. Minimizing Eq. S1 again with respect to $\gamma$, we now get:

$$\frac{(R-h)^2}{R^2\left(1-\gamma_{\text{cyl}}\right)^2} - \frac{(R+h)^2}{R^2\left(1+\gamma_{\text{cyl}}\right)^2} = \frac{\kappa}{\mu_A}\left[\left(\frac{1}{R-h} - c_0\right)^2 - \left(\frac{1}{R+h} + c_0\right)^2\right] \quad (S4)$$

To simplify, the right-hand side of Eq. 8 is zero if: (i) $c_0 = h/(R^2 - h^2)$; or (ii) $R$ is large, resulting in:

$$\gamma_{\text{cyl}} \approx \frac{h}{R} \tag{S5}$$

Similar to the sphere case, the monolayer equilibrium area is now independent of material properties, and we correctly recover that as the curvature flattens, $R \to +\infty$ and $\gamma_{\text{cyl}} \to 0$. In the numerical simulations, we use the exact values numerically solved from Eq. S2 and Eq. S4.

#### II. Height energy function that accounts for membrane thinning and thickening

Our theory in the manuscript shows that the membrane tilt energy can be reformulated as a height energy function per unit area,

$$\bar{E}_h^{\text{thin}} = \frac{1}{2}\kappa_h\left(\rho_{app} - \frac{1}{3}\right)^2 - \frac{\kappa_h}{18}, \quad (-1 \le \rho_{app} < 0) \tag{SI 1}$$

where $\kappa_h$ is the height modulus, $\rho_{app} = (h - h_0^C)/h_0^C$ is the height strain, $h$ is the observed monolayer height, and $h_0^C$ is the curvature-modified height, which is also the equilibrium height. The above function applies only to the membrane thinning ($-1 \le \rho_{app} < 0$ or $0 < h \le h_0^c$), as tilt can only cause membrane thinning. However, the energy minimization of this equation will drive $h$ to converge to $4h_0^C/3$.

To address this, we introduce a constraint energy that penalizes cases where $h > h_0^c$:

$$\bar{E}_h^{\text{pen}} = \frac{1}{2}\mu_h'\rho_{app}^2, \quad (\rho_{app} > 0). \tag{SI 2}$$

The artificial coefficient $\mu_h'$ is set large enough that this term becomes negligible in equilibrium, effectively acting as a strict constraint against membrane thickening. Without loss of generality, we set $\mu_h' = 1GPa \times 2.5nm = 2.5 \times 10^3\ pN/nm$, where $1GPa$ is the membrane's bulk modulus, and 2.5 nm is the typical thickness of the monolayer.

It is important to note that the inclusion of Eq(S2) to the energy function in Eq(S1) ensures the function value is continuous with respect to $\rho_{app}$ (or $h$) over the entire range of values, but the first derivative is not continuous at $\rho_{app} = 0$ (or $h = h_0^C$). This non-differentiability reduces the efficiency of energy minimization. To resolve this issue, we develop the following piecewise functions for the height energy,

$$\bar{E}_h = \begin{cases} \frac{1}{2}\kappa_h\left(\rho_{app} - \frac{1}{3}\right)^2 - \frac{1}{18}\kappa_h, & -1 < \rho_{app} \le -\varepsilon \\ A\rho_{app}^3 + B\rho_{app}^2, & -\varepsilon < \rho_{app} \le 0 \\ \frac{1}{2}\mu_h\rho_{app}^2, & \rho_{app} > 0 \end{cases} \tag{SI 3}$$

Here, the coefficients A and B are selected to ensure the function is differentiable across the entire domain:

$$\begin{cases} A = -\frac{\kappa_h}{\varepsilon^3}\left(\varepsilon - \frac{1}{3}\right)^2 + \frac{\kappa_h}{\varepsilon^2}\left(\varepsilon - \frac{1}{3}\right) + \frac{\kappa_h}{9\varepsilon^3} \\ B = \frac{3\kappa_h}{2\varepsilon^2}\left(\varepsilon - \frac{1}{3}\right)^2 - \frac{\kappa_h}{\varepsilon}\left(\varepsilon - \frac{1}{3}\right) - \frac{\kappa_h}{6\varepsilon^2} \end{cases},$$

with $\varepsilon = 10^{-2}$, a small arbitrary value. The second piecewise function in Eq(S3) applies only to a small region around $\rho_{app} = 0$, so its influence on the energy minimum is negligible. By incorporating this function into the energy framework, both the function value and its first derivative become continuous with respect to $\rho_{\text{app}}$ across the entire domain.

### III. Kinematics

**Background on differential geometry**

We model the lipid membrane as two coupled surfaces, each of which is parameterized by $\vec{x}(s^1, s^2)$, where $\{s^1, s^2\}$ are curvilinear coordinates. Details on the geometry of surfaces can be found in [3, 4]. Here, we just provide the fundamental mathematics and relevant derivative functions that we used in this work [5]. The covariant basis vector $\vec{a}_\beta$ at each surface point is defined as

$$\vec{a}_\beta = \frac{\partial \vec{x}}{\partial s^\beta},$$

where $\beta$ = 1, 2. The unit normal to the surface is

$$\vec{d} \equiv \vec{a}_3 = \frac{\vec{a}_1 \times \vec{a}_2}{\sqrt{a}},$$

where $\sqrt{a} = |\vec{a}_1 \times \vec{a}_2|$. The infinitesimal element of the surface area at the point $\vec{x}(s^1, s^2)$ is

$$dA = \sqrt{a} ds^1 ds^2 \equiv \sqrt{a} d^2 s.$$

The contravariant basis vector $\vec{a}^\beta$ is defined such that $\vec{a}^\beta \cdot \vec{a}_\alpha = \delta_\alpha^\beta$, or

$$\vec{a}^1 = \frac{\vec{a}_2 \times \vec{a}_3}{\sqrt{a}}, \text{ and } \vec{a}^2 = \frac{\vec{a}_3 \times \vec{a}_1}{\sqrt{a}}.$$

The covariant and contravariant components of the surface metric tensor follow as

$$a_{\alpha\beta} = \vec{a}_\alpha \cdot \vec{a}_\beta, \text{ and } a^{\alpha\beta} = \vec{a}^\alpha \cdot \vec{a}^\beta.$$

Here, the Greek indices $\alpha$ and $\beta$ both take values of 1 and 2. The local curvature measures the change in the normal and the curvature tensor is

$$B_{\alpha\beta} = \vec{d} \cdot \frac{\partial \vec{a}_\alpha}{\partial s^\beta}.$$

The mean curvature is half of the trace of the curvature tensor,

$$c = \frac{1}{2} tr(B) \equiv B_{\alpha\alpha}.$$

It is noted that the Einstein summation convention is used from now on.
The Gaussian curvature is the determinant of the curvature tensor,

$$c_G = det(B).$$

**Application to the finite element method**

We use the triangular mesh for the finite element analysis of the membrane [6]. The membrane surface is the limit surface obtained from the subdivision of triangulations with Loop's scheme [7]. Consequently, the local membrane over the triangular element *e* can be parameterized by

$$\vec{x}(s^1, s^2) = \sum_{i=1}^{12} \vec{x}_i N_i(s^1, s^2),$$

where $\{s^1, s^2\}$ is chosen as two of barycentric coordinates in triangle *e*, $\vec{x}_i$ is the nodal position of the one-ring neighbor vertices around the triangle *e*, and $N_i(s^1, s^2)$ is the box-spline shape function [7]. In this finite element mesh, the basis vectors of the surface are calculated as the linear combination of one-ring vertices,

$\vec{a}_\beta = \frac{\partial \vec{x}}{\partial s^\beta} = \sum_{i=1}^{12} \vec{x}_i \frac{\partial N_i}{\partial s^\beta}$ .

The derivative of the basis vector to the barycentric coordinate is also the linear combination of one-ring vertices,

$\frac{\partial \vec{a}_\alpha}{\partial s^\beta} = \sum_{i=1}^{12} \vec{x}_i \frac{\partial^2 N_i}{\partial s^\alpha \partial s^\beta}$.

By the definition of $\vec{a}_x = \vec{a}_1 \times \vec{a}_2$, we have

$\frac{\partial \vec{a}_x}{\partial s^\beta} = \left(\frac{\partial \vec{a}_1}{\partial s^\beta}\right) \times \vec{a}_2 + \vec{a}_1 \times \left(\frac{\partial \vec{a}_2}{\partial s^\beta}\right)$ .

$\frac{\partial \sqrt{a}}{\partial s^\beta} = \frac{1}{\sqrt{a}} \vec{a}_x \cdot \frac{\partial \vec{a}_x}{\partial s^\beta}$ .

$\frac{\partial \vec{d}}{\partial s^\beta} = \frac{1}{\left(\sqrt{a}\right)^2} \left(\frac{\partial \vec{a}_x}{\partial s^\beta} \sqrt{a} + \vec{a}_x \cdot \frac{\partial \sqrt{a}}{\partial s^\beta}\right)$.

$\frac{\partial \vec{a}^1}{\partial s^\beta} = \frac{1}{\left(\sqrt{a}\right)^2} \left[\left(\frac{\partial \vec{a}_2}{\partial s^\beta} \times \vec{d} + \vec{a}_2 \times \frac{\partial \vec{d}}{\partial s^\beta}\right) \sqrt{a} - \left(\vec{a}_2 \times \vec{d}\right) \frac{\partial \sqrt{a}}{\partial s^\beta}\right]$ .

$\frac{\partial \vec{a}^2}{\partial s^\beta} = \frac{1}{\left(\sqrt{a}\right)^2} \left[\left(\frac{\partial \vec{d}}{\partial s^\beta} \times \vec{a}_1 + \vec{d} \times \frac{\partial \vec{a}_1}{\partial s^\beta}\right) \sqrt{a} - \left(\vec{d} \times \vec{a}_1\right) \frac{\partial \sqrt{a}}{\partial s^\beta}\right]$ .

It is also practical to calculate the derivative to the nodal vertex position, such as

$\frac{\partial \vec{x}}{\partial \vec{x}_i} = N_i[I]$,

$\frac{\partial \vec{a}_\beta}{\partial \vec{x}_i} = \frac{\partial N_i}{\partial s^\beta}[I]$,

$\frac{\partial \sqrt{a}}{\partial \vec{x}_i} = \sqrt{a}\left(\vec{a}^\beta \cdot \frac{\partial \vec{a}_\beta}{\partial \vec{x}_i}\right)$,

$\frac{\partial \vec{d}}{\partial \vec{x}_i} = -\frac{\partial N_i}{\partial s^\beta}\left[\left(\vec{a}^\beta\right)^T \otimes \vec{d}\right]$,

$\frac{\partial}{\partial \vec{x}_i}\left(\frac{\partial \vec{d}}{\partial s^\beta}\right) = \frac{\partial}{\partial s^\beta}\left(\frac{\partial \vec{d}}{\partial \vec{x}_i}\right) = -\frac{\partial^2 N_i}{\partial s^\beta \partial s^\alpha}\left[(\vec{a}^\alpha)^T \otimes \vec{d}\right] - \frac{\partial N_i}{\partial s^\alpha}\left[\left(\frac{\partial \vec{a}^\alpha}{\partial s^\beta}\right)^T \otimes \vec{d}\right] - \frac{\partial N_i}{\partial s^\alpha}\left[(\vec{a}^\alpha)^T \otimes \frac{\partial \vec{d}}{\partial s^\beta}\right]$ ,

where $[I]$ is 3×3 identity matrix, $T$ is the transpose symbol and ⊗ is the tensor product symbol. It is noted that we use row vectors during our calculation, and the tensor product here is performed between a column vector and a row vector. The derivative of the total curvature to the vertex position is

$\frac{\partial c}{\partial \vec{x}_i} = -\frac{1}{2}\left[\vec{a}^\beta \cdot \frac{\partial}{\partial \vec{x}_i}\left(\frac{\partial \vec{d}}{\partial s^\beta}\right)\right] + \frac{1}{2}\left[\left(\vec{a}^\alpha \cdot \vec{a}^\beta\right) \frac{\partial \vec{d}}{\partial s^\alpha} \cdot \frac{\partial \vec{a}_\beta}{\partial \vec{x}_i}\right]$.

#### IV. Finite element analysis of the membrane mechanics

Our bilayer membrane model includes the bending energetic $E_c$ and area energetic $E_A$ of the two monolayers, both of which have similar definition as the previous one-layer surface in the curvilinear coordinate framework [5]. The inner and outer monolayers follow the same definitions. The energy $E_V$ of the constraint on the volume that the membrane encloses, such as the vesicle, has also been previously defined [5]. We will derive the new energy term due to the monolayer height deformation of the double-layer membrane system, $E_h$.

**Membrane energetics**

The total energy of our bilayer membrane model is

$$E_{tot} = (E_c^I + E_A^I + E_h^I) + (E_c^O + E_A^O + E_h^O) + E_V \ . \tag{SI 4}$$

where the upper subscript '$I$' and '$O$' refer to the inner and outer monolayer, respectively. For the planar membrane, '$I$' and '$O$' will refer to the bottom and top monolayer, respectively. In the following, we will express each energetic term in the curvilinear coordinates $\{s^1, s^2\}$.

*1. Previously defined terms*

For each monolayer, the bending energy follows the well-known Helfrich Hamiltonian,

$$\begin{cases} E_c^I = \int \frac{1}{2}\kappa^I (2c^I - c_0^I)^2 \sqrt{a}^I \, d^2s \\ E_c^o = \int \frac{1}{2}\kappa^o (2c^o - c_0^o)^2 \sqrt{a}^o \, d^2s \end{cases}, \tag{SI 5}$$

where the parameters: $\kappa$ is the monolayer bending modulus, $c_0$ is the monolayer spontaneous curvature or lipid spontaneous curvature; the variables: $c$ is the mean curvature. $\sqrt{a}d^2s \equiv \sqrt{a}ds^1ds^2$ is the infinitesimal element of the area, see the Kinematic section. The integral is performed over the monolayer area.

The area elasticity of each monolayer describes the stretch of the global area,

$$\begin{cases} E_A^I = \frac{1}{2}\mu_A^I \frac{\left(A^I - A_0^I\right)^2}{A_0^I} \\ E_A^o = \frac{1}{2}\mu_A^o \frac{(A^o - A_0^o)^2}{A_0^o} \end{cases}, \tag{SI 6}$$

where the parameters: $\mu_A$ is the area elasticity modulus of the monolayer, $A_0$ is the monolayer equilibrium area; the variable $A = \int \sqrt{a}\, d^2s$ is the global area of the monolayer.

If the membrane is an enclosed vesicle, there is an energetic for the vesicle volume constraint, which is expressed by

$$E_V = \frac{1}{2}\mu_V \frac{(V - V_0)^2}{V_0}, \tag{SI 7a}$$

where the parameters $\mu_V$ is the constant of the volume constraint, and $V_0$ is the equilibrium value of the vesicle volume. The variable $V$ is the volume enclosed by the inner monolayer of the bilayer membrane,

$$V = V^I = \int \frac{1}{3}\left(\vec{x}^I \cdot \vec{d}^I\right)\sqrt{a}^I \, d^2s, \tag{SI 7b}$$

where $\vec{x}^I = \vec{x}^I(s^1, s^2)$ is each point on the inner monolayer surface, $\vec{d}^I$ is the surface normal at the point $\vec{x}^I$, and $\sqrt{a}^I d^2s$ is the infinitesimal area at the point $\vec{x}^I$.

*2. New terms*

We define the height deformation of the bilayer membrane in an energetic term,

$$E_h^I = \begin{cases} \int \left[\frac{1}{2}\kappa_h^I \left(\rho^I - \frac{1}{3}\right)^2 - \frac{1}{18}\kappa_h^I\right] \sqrt{a}^I \, d^2s, & (-1 < \rho^I \le -\varepsilon) \\ \int \left[A{\rho^I}^3 + B{\rho^I}^2\right] \sqrt{a}^I \, d^2s, & (-\varepsilon < \rho^I \le 0) \\ \int \frac{1}{2}\mu_h^I {\rho^I}^2 \sqrt{a}^I \, d^2s, & (0 < \rho^I) \end{cases} \quad \text{(SI 8a)}$$

$$E_h^O = \begin{cases} \int \left[\frac{1}{2}\kappa_h^O \left(\rho^O - \frac{1}{3}\right)^2 - \frac{1}{18}\kappa_h^O\right] \sqrt{a}^I \, d^2s, & (-1 < \rho^O \le -\varepsilon) \\ \int \left[A{\rho^O}^3 + B{\rho^O}^2\right] \sqrt{a}^O \, d^2s, & (-\varepsilon < \rho^O \le 0) \\ \int \frac{1}{2}\mu_h^O {\rho^O}^2 \sqrt{a}^O \, d^2s, & (0 < \rho^O) \end{cases} \quad \text{(SI 8b)}$$

where the parameters: $\kappa_h^I$, $\kappa_h^O$ are the height modulus of the inner and outer monolayer, respectively; $\rho^I = \frac{h^I - h_0^{cI}}{h_0^{cI}}$, and $h_0^{cI}$ is the curvature-modified equilibrium height of the inner monolayer; $\rho^O = \frac{h^O - h_0^{cO}}{h_0^{cO}}$, and $h_0^{cO}$ is the curvature-modified equilibrium height of the outer monolayer. Note $h_0^{cI}$ and $h_0^{cO}$ are determined at the beginning of the simulations, while during the simulation their value are fixed. The integral of Eq (SI 8) is performed over the monolayer area. The variable $h$ is the observed height of the monolayer, which is defined by

$$\begin{cases} h^I = [\vec{x}^I(s^1, s^2) - \vec{x}^M(s^1, s^2)] \cdot \vec{d}^I \\ h^O = [\vec{x}^o(s^1, s^2) - \vec{x}^M(s^1, s^2)] \cdot \vec{d}^o \end{cases}, \quad \text{(SI 9)}$$

where $\vec{x}^I$, $\vec{x}^M$ and $\vec{x}^O$ are three points on the inner, middle and outer surface, respectively. The vector $\vec{d}^I$ is the normal to the inner monolayer surface at the point $\vec{x}^I$, and $\vec{d}^o$ is the normal to the outer monolayer surface at the point $\vec{x}^o$. It is noted that our bilayer membrane model is built by three layers of triangular mesh, i.e., the inner, middle and outer layers use the same finite element indexes and vertex indexes. The three points, $\vec{x}^I$, $\vec{x}^M$ and $\vec{x}^O$, in Eq (SI 9) are chosen with the same curvilinear coordinates $(s^1, s^2)$. Without deformation, the distance between $\vec{x}^o$ and $\vec{x}^I$ is equal to the bilayer thickness, and the line connected by them is parallel to the surface normal $\vec{d}^o$.

**Nodal force**

The membrane deformation will generate forces on its surfaces. Here, we use triangular finite elements to simulate the membrane. The force $\vec{F}_i$ imposed on the mesh vertex $\vec{x}_i$ can be calculated by the derivative of the total energy to the vertex position, $\vec{F}_i = -\frac{\partial E_{tot}}{\partial \vec{x}_i}$. In the following context, we will provide the detailed expression of the nodal force on the outer, inner and middle mesh, respectively.

*1. Nodal force on the outer mesh layer*

Consider a vertex on the outer triangular mesh, $\vec{x}_i^o$. The energetic terms that involve the outer vertex include the bending energy of the outer monolayer $E_c^o$, Eq (SI 5), the area elasticity of the outer monolayer $E_A^o$, Eq (SI 6), and the monolayer height elasticity $E_h^o$, Eq (SI 8). The nodal force imposed on the vertex $\vec{x}_i^o$ should cover all these three energetic effects,

$$\vec{F}_i^o = -\frac{\partial E_{tot}}{\partial \vec{x}_i^o} = -\frac{\partial E_c^o}{\partial \vec{x}_i^o} - \frac{\partial E_A^o}{\partial \vec{x}_i^o} - \frac{\partial E_h^o}{\partial \vec{x}_i^o}. \quad \text{(SI 10a)}$$

The force component raised from the curvature change is

$$\vec{F}_{c,i}^o = -\frac{\partial E_c^o}{\partial \vec{x}_i^o} = -\frac{\kappa^o}{2}\int\left[4(2c^o - c_0^o)\sqrt{a}^o\frac{\partial c^o}{\partial \vec{x}_i^o} + (2c^o - c_0^o)^2\frac{\partial \sqrt{a}^o}{\partial \vec{x}_i^o}\right]d^2s, \quad \text{(SI 10b)}$$

where the detailed expression of $\frac{\partial c^o}{\partial \vec{x}_i^o}$ and $\frac{\partial \sqrt{a}^o}{\partial \vec{x}_i^o}$ can be found in the Kinematics section.

The force component raised from the area change is

$$\vec{F}_{A,i}^o = -\frac{\partial E_A^o}{\partial \vec{x}_i^o} = -\frac{\mu_A^o}{A_0^o}(A^o - A_0^o)\int\frac{\partial \sqrt{a}^o}{\partial \vec{x}_i^o}d^2s, \quad \text{(SI 10c)}$$

where $\frac{\partial \sqrt{a}^o}{\partial \vec{x}_i^o}$ is expressed in the Kinematics section.

The force component raised from the height change is,

$$\vec{F}_{h,i}^o = -\frac{\partial E_h^o}{\partial \vec{x}_i^o}$$

$$= \begin{cases} -\int\left\{\kappa_h^O\left(\rho^O - \frac{1}{3}\right)\frac{\partial \rho^O}{\partial \vec{x}_i^O}\sqrt{a}^O + \left[\frac{\kappa_h^O}{2}\left(\rho^O - \frac{1}{3}\right)^2 - \frac{\kappa_h^O}{18}\right]\frac{\partial \sqrt{a}^O}{\partial \vec{x}_i^O}\right\}d^2s, (-1 < \rho^O \le -\varepsilon) \\ -\int\left\{\left(3A{\rho^O}^2 + 2B\rho^O\right)\frac{\partial \rho^O}{\partial \vec{x}_i^O}\sqrt{a}^O + \left[A{\rho^O}^3 + B{\rho^O}^2\right]\frac{\partial \sqrt{a}^O}{\partial \vec{x}_i^O}\right\}d^2s, \ (-\varepsilon < \rho^O \le 0) \\ -\int\left\{\mu_h^O\rho^O\frac{\partial \rho^O}{\partial \vec{x}_i^O}\sqrt{a}^O + \frac{1}{2}\mu_h^O{\rho^O}^2\frac{\partial \sqrt{a}^O}{\partial \vec{x}_i^O}\right\}d^2s, \qquad (0 < \rho^O) \end{cases} \quad \text{(SI 10d)}$$

where $\frac{\partial \rho^O}{\partial \vec{x}_i^O} = \frac{1}{h_0^{cO}}\frac{\partial h^O}{\partial \vec{x}_i^O}$, and $\frac{\partial h^O}{\partial \vec{x}_i^O} = \frac{\partial \vec{x}^o}{\partial \vec{x}_i^O}\cdot\vec{d}^o + (\vec{x}^o - \vec{x}^M)\cdot\frac{\partial \vec{d}^o}{\partial \vec{x}_i^O}$ ; $\vec{x}^o$, $\vec{x}^M$ are two points on outer and mid-surface mesh, respectively, defining the monolayer height. The expression of $\frac{\partial \vec{x}^o}{\partial \vec{x}_i^O}$, $\frac{\partial \vec{d}^o}{\partial \vec{x}_i^O}$, $\frac{\partial \sqrt{a}^O}{\partial \vec{x}_i^O}$ and $\frac{\partial c^O}{\vec{x}_i^O}$ can be found in the Kinematics section.

*2. Nodal force on the inner mesh layer*

Consider a vertex on the inner triangular mesh, $\vec{x}_i^I$. The energetic terms that involve the inner vertex include the bending energy of the inner monolayer $E_c^I$, Eq (SI 5), the area elasticity of the inner monolayer $E_A^I$, Eq (SI 6), the monolayer height elasticity $E_h^I$, Eq (SI 8), and the volume constraint energy $E_V$, Eq (SI 7). The nodal force imposed on the vertex $\vec{x}_i^I$ should cover all these four energetic effects,

$$\vec{F}_i^I = -\frac{\partial E_{tot}}{\partial \vec{x}_i^I} = -\frac{\partial E_c^I}{\partial \vec{x}_i^I} - \frac{\partial E_A^I}{\partial \vec{x}_i^I} - \frac{\partial E_h^I}{\partial \vec{x}_i^I} - \frac{\partial E_V}{\partial \vec{x}_i^I}. \quad \text{(SI 11a)}$$

The force component raised from the curvature change is

$$\vec{F}_{c,i}^I = -\frac{\partial E_c^I}{\partial \vec{x}_i^I} = -\frac{\kappa^I}{2}\int\left[4(2c^I - c_0^I)\sqrt{a}^I\frac{\partial c^I}{\partial \vec{x}_i^I} + (2c^I - c_0^I)^2\frac{\partial \sqrt{a}^I}{\partial \vec{x}_i^I}\right]d^2s, \quad \text{(SI 11b)}$$

where the detailed expression of $\frac{\partial c^I}{\partial \vec{x}_i^I}$ and $\frac{\partial \sqrt{a}^I}{\partial \vec{x}_i^I}$ can be found in the Kinematics section.

The force component raised from the area change is

$$\vec{F}_{A,i}^I = -\frac{\partial E_A^I}{\partial \vec{x}_i^I} = -\frac{\mu_A^I}{A_0^I}(A^I - A_0^I)\int\frac{\partial \sqrt{a}^I}{\partial \vec{x}_i^I}d^2s, \quad \text{(SI 11c)}$$

where $\frac{\partial \sqrt{a}^I}{\partial \vec{x}_i^I}$ is expressed in the Kinematics section.

The force component raised from the inner height change is,

$$\vec{F}_{h,i}^I = -\frac{\partial E_h^I}{\partial \vec{x}_i^I}$$

$$= \begin{cases} -\int \left\{ \kappa_h^I \left(\rho^I - \frac{1}{3}\right) \frac{\partial \rho^I}{\partial \vec{x}_i^I} \sqrt{a}^I + \left[ \frac{\kappa_h^I}{2}\left(\rho^I - \frac{1}{3}\right)^2 - \frac{\kappa_h^I}{18} \right] \frac{\partial \sqrt{a}^I}{\partial \vec{x}_i^I} \right\} d^2 s, (-1 < \rho^I \le -\varepsilon) \\ -\int \left\{ \left(3A{\rho^I}^2 + 2B\rho^I\right) \frac{\partial \rho^I}{\partial \vec{x}_i^I} \sqrt{a}^I + \left[ A{\rho^I}^3 + B{\rho^I}^2 \right] \frac{\partial \sqrt{a}^I}{\partial \vec{x}_i^I} \right\} d^2 s, \ (-\varepsilon < \rho^I \le 0) \\ -\int \left\{ \mu_h^I \rho^I \frac{\partial \rho^I}{\partial \vec{x}_i^I} \sqrt{a}^I + \frac{1}{2} \mu_h^I {\rho^I}^2 \frac{\partial \sqrt{a}^I}{\partial \vec{x}_i^I} \right\} d^2 s, \quad (0 < \rho^I) \end{cases} \quad \text{(SI 11d)}$$

where $\frac{\partial \rho^I}{\partial \vec{x}_i^I} = \frac{1}{h_0^{cI}} \frac{\partial h^I}{\partial \vec{x}_i^I}$, and $\frac{\partial h^I}{\partial \vec{x}_i^I} = \frac{\partial \vec{x}^I}{\partial \vec{x}_i^I} \cdot \vec{d}^I + (\vec{x}^I - \vec{x}^M) \cdot \frac{\partial \vec{d}^I}{\partial \vec{x}_i^I}$, and $\vec{x}^I$, $\vec{x}^M$ are two vertices on inner and mid-surface mesh, respectively, defining the monolayer height. The expression of $\frac{\partial \vec{x}^I}{\partial \vec{x}_i^I}$, $\frac{\partial \vec{d}^I}{\partial \vec{x}_i^I}$, $\frac{\partial \sqrt{a}^I}{\partial \vec{x}_i^I}$ and $\frac{\partial c^I}{\vec{x}_i^I}$ can be found in the Kinematics section.

The force component raised from the change of the vesicle volume is

$$\vec{F}_{V,i}^I = -\frac{\partial E_V}{\partial \vec{x}_i^I}$$

$$= -\frac{1}{3}\frac{\mu_V}{V_0^I}(V^I - V_0^I) \int \left[ \frac{\partial \vec{x}^I}{\partial \vec{x}_i^I} \cdot \vec{d}^I \sqrt{a}^I + \vec{x}^I \cdot \frac{\partial \vec{d}^I}{\partial \vec{x}_i^I} \sqrt{a}^I + \vec{x}^I \cdot \vec{d}^I \cdot \frac{\partial \sqrt{a}^I}{\partial \vec{x}_i^I} \right] d^2 s \ , \quad \text{(SI 11f)}$$

where the detailed expressions of $\frac{\partial \vec{x}^I}{\partial \vec{x}_i^I}$, $\frac{\partial \vec{d}^I}{\partial \vec{x}_i^I}$, and $\frac{\partial \sqrt{a}^I}{\partial \vec{x}_i^I}$ can be found in the Kinematics section.

*3. Nodal force on the mid-surface layer*

Consider a vertex on the midplane triangular mesh, $\vec{x}_i^M$. The energetic terms that involve the midplane vertex include the monolayer height elasticity $E_h^O$ and $E_h^I$, Eq (SI 8). The nodal force imposed on the vertex $\vec{x}_i^M$ should cover all these two energetic effects,

$$\vec{F}_i^M = -\frac{\partial E_{tot}}{\partial \vec{x}_i^M} = -\frac{\partial E_h^O}{\partial \vec{x}_i^M} - \frac{\partial E_h^I}{\partial \vec{x}_i^M}, \quad \text{(SI 12)}$$

where,

$$\frac{\partial E_h^O}{\partial \vec{x}_i^M} = \begin{cases} \int \left\{ \kappa_h^O \left(\rho^O - \frac{1}{3}\right) \frac{\sqrt{a}^O}{h_0^{cO}} \vec{d}^O \right\} d^2 s, \quad (-1 < \rho^O \le -\varepsilon) \\ \int \left\{ \left(3A{\rho^O}^2 + 2B\rho^O\right) \frac{\sqrt{a}^O}{h_0^{cO}} \vec{d}^O \right\} d^2 s, \ (-\varepsilon < \rho^O \le 0) \\ \int \left\{ \mu_h^O \rho^O \frac{\sqrt{a}^O}{h_0^{cO}} \vec{d}^O \right\} d^2 s, \quad (0 < \rho^O) \end{cases}$$

and

$$\frac{\partial E_h^I}{\partial \vec{x}_i^M} = \begin{cases} \int \left\{ \kappa_h^I \left(\rho^I - \frac{1}{3}\right) \frac{\sqrt{a}^I}{h_0^{cI}} \vec{d}^I \right\} d^2 s, \quad (-1 < \rho^I \le -\varepsilon) \\ \int \left\{ \left(3A{\rho^I}^2 + 2B\rho^I\right) \frac{\sqrt{a}^I}{h_0^{cI}} \vec{d}^I \right\} d^2 s, \ (-\varepsilon < \rho^I \le 0) \\ \int \left\{ \mu_h^I \rho^I \frac{\sqrt{a}^I}{h_0^{cI}} \vec{d}^I \right\} d^2 s, \quad (0 < \rho^I) \end{cases}$$

**V. Increasing mesh resolution around the insertion in the continuum membrane**

The helix insertion on the membrane generates a membrane deformation distributed relatively locally around the insertion. To describe the local deformation more precisely in the finite element method, we use a finer mesh. Because a finer mesh is more computationally expensive, we only apply it close to the insertion, reverting to a coarser mesh at larger distances (Fig S1). This locally finer mesh includes regular patches (triangles) and three types of irregular patches (see Table S1). For irregular patches, we apply the subdivision method [5] to generate 4 smaller patches, and any remaining irregular patches are subdivided once more.

The area of the finer mesh is defined to be large enough to incorporate the local area perturbed by the helix insertion (Fig S1). We find that this area is less than 4 nm in radius from the insertion, as we show below. Therefore, we set the local area of the finer mesh to a radius of 5 nm. We note that the membrane energy does decrease slightly as the mesh becomes finer around the insertion (FigS1). Though making the mesh infinitely finer around the insertion improves the mathematical precision of the calculation, this is not physically realistic. Each lipid has an area ~0.5 $nm^2$ in average, which suggests a lower length limit of ~0.7 nm for a rigid deformation distance on the mesh, otherwise the surface structure is capturing structural variation below the molecular scale. We set the smallest triangle edge as 0.5 nm as a comparable lower bound for the finer mesh.

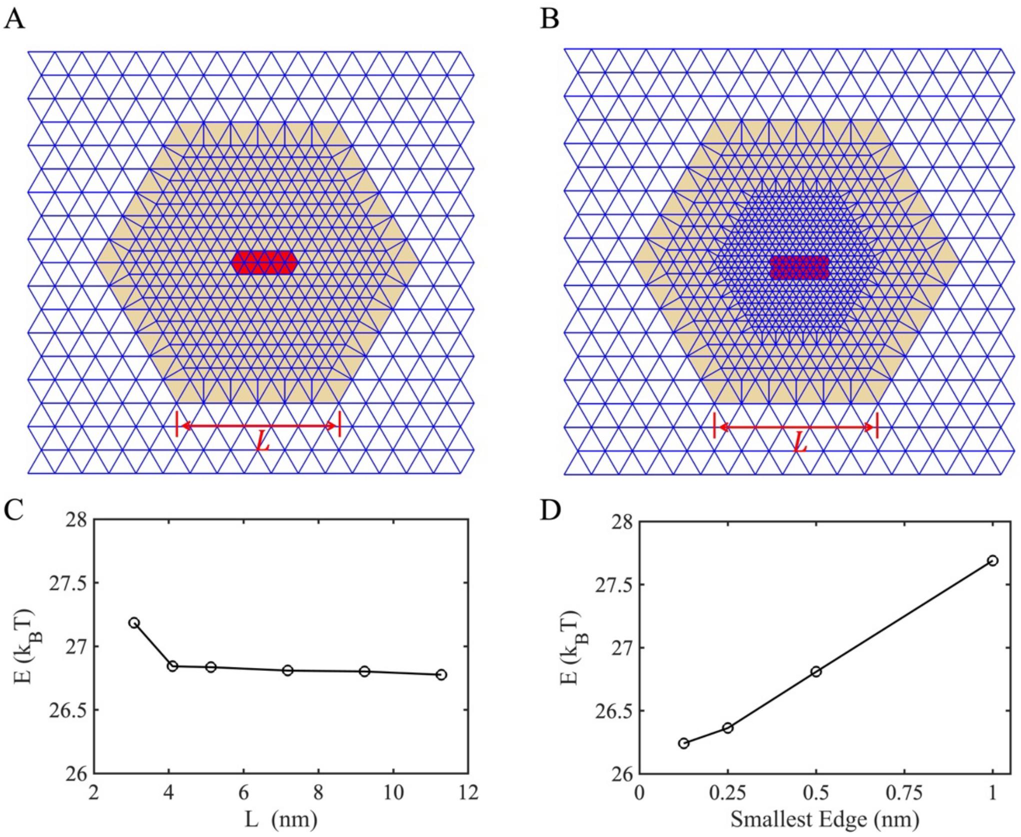


Figure S1. Locally finer mesh. (A) A zone on the mesh (orange color) is selected to build the finer mesh through triangular subdivisions. The red-color zone is the location for helix insertion. (B) Similar to (A) but the mesh around the insertion is furtherly finer. (C) The size of the area of the finer mesh doesn't affect the energy calculation as long as $L \geq 4$ nm. (D) The calculated energy of the membrane decreases as the local mesh becomes finer and finer.

**Table S1. Five types of patches on the triangular mesh**

| Patch Type | Patch Sample | Characters | Subdivision | Sub-patch Type |
|---|---|---|---|---|
| Regular | 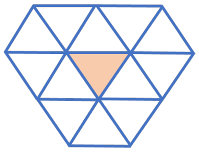 | 12 one-ring vertices<br>valences = [6, 6, 6] | — | — |
| Irregular-minus | 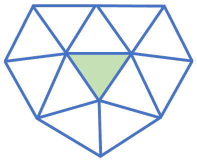 | 11 one-ring vertices<br>valences = [5, 6, 6] | 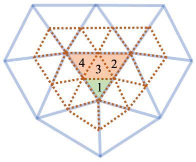 | 1: irregular-minus<br>2: regular<br>3: regular<br>4: regular |
| Irregular-plus | 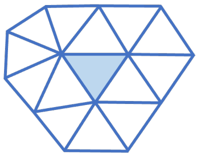 | 13 one-ring vertices<br>valences = [6, 6, 7] | 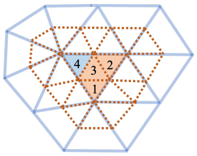 | 1: regular<br>2: regular<br>3: regular<br>4: irregular-plus |
| Pseudo-regular | 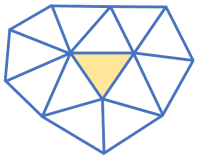 | 12 one-ring vertices<br>valences = [5, 6, 7] | 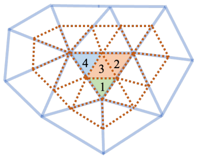 | 1: irregular-minus<br>2: regular<br>3: regular<br>4: irregular-plus |

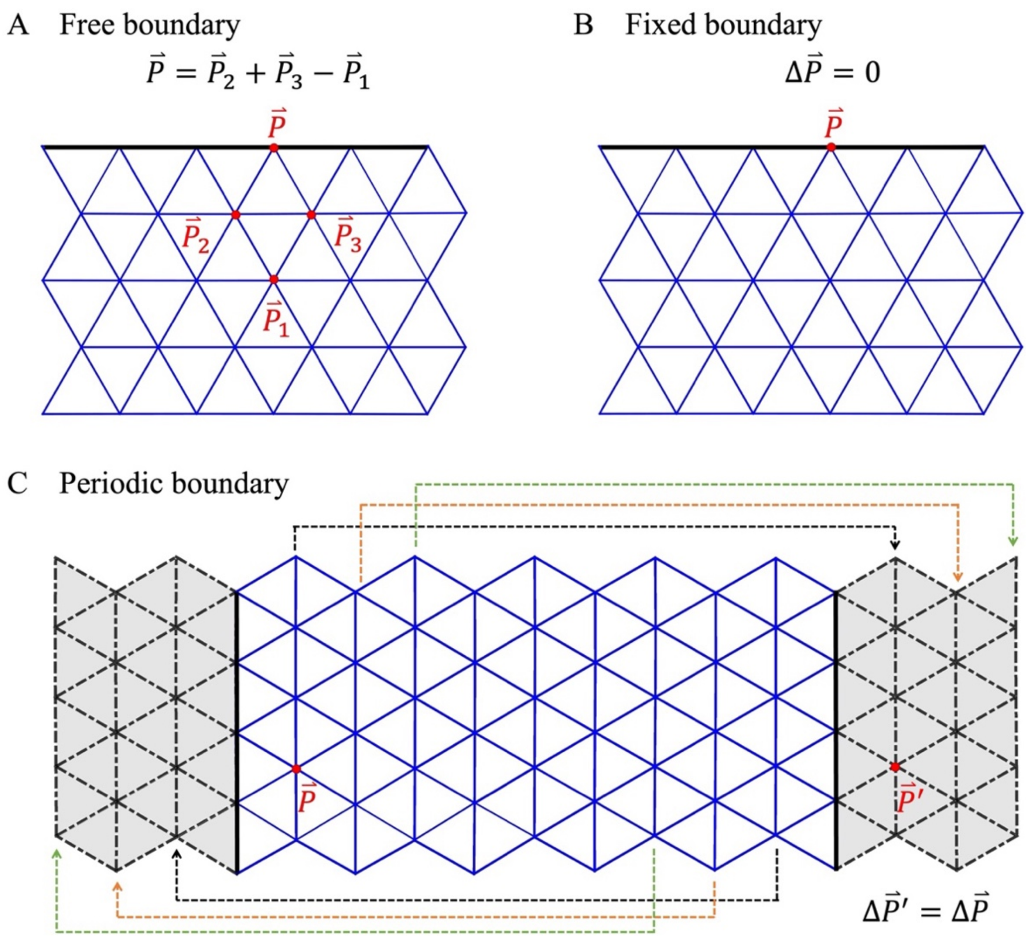


**Figure S2. Schematics of three boundary conditions for planar or cylindrical membranes.** The blue triangular meshes represent the membrane region. The black solid lines represent the boundary lines or edges of the membrane. The black dashed lines or gray areas represent the ghost zones. (A) For the free boundary condition, the position of vertex $\vec{P}$ on the membrane edge is determined by its three neighboring vertices, $\vec{P}_1$, $\vec{P}_2$ and $\vec{P}_3$. (B) For the fixed boundary condition, the vertex on the edge remains immovable throughout the simulation, so the vertex displacement $\Delta\vec{P}$ is 0. (C) For the periodic boundary condition, ghost vertices and faces (gray color) are added to each edge of the membrane, which aims to help the calculation of the differential geometry on the real vertices and faces near the edge. The position of the ghost vertex is updated by the translation of the real vertex position on the relatively opposite end of the membrane, as shown by the colored dashed lines. For example, the position of the ghost vertex $\vec{P}'$ on the right is obtained by the translation of the real vertex $\vec{P}$ on the left, so their vertex displacements are equivalent to each other, $\Delta\vec{P}' = \Delta\vec{P}$.

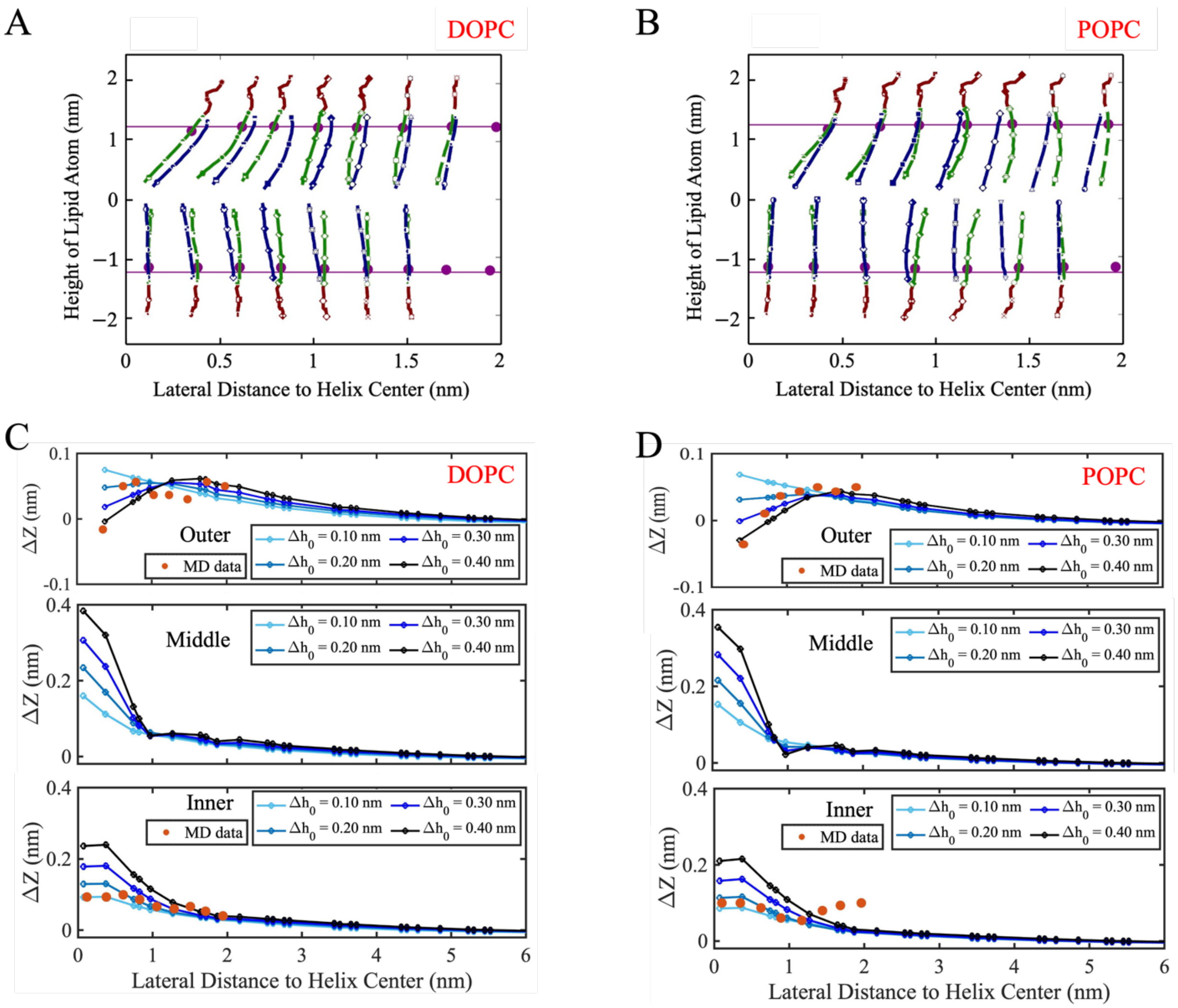


**Figure S3. Membrane deformations observed through MD simulations and the continuum membrane model.** MD simulations depict lipid tilting and leaflet thinning around the helix insertion in membranes with DOPC lipids (A) and POPC lipids (B). The continuum membrane model similarly captures the deformations of the outer leaflet, inner leaflet and the middle surface caused by the helix insertion in DOPC (C) and POPC (D) membranes. The red dots in (C) and (D) correspond to the dark purple dots in (A) and (B), respectively.

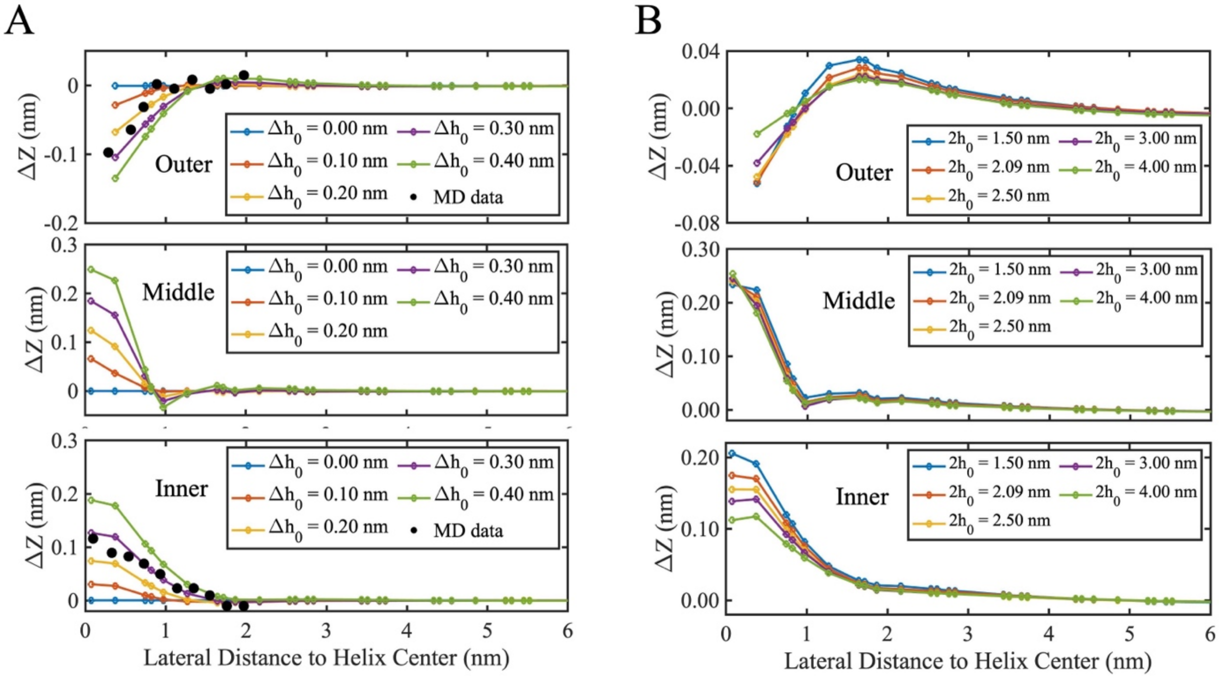


**Figure S4. The DLPC lipid membrane is deformed due to the helix insertion, demonstrated by our continuum membrane model.** (A) The height deformation of the membrane changes with the distance from the helix insertion, depending on $\Delta h_0$, where $c_{0,ins} = 0.1$ nm$^{-1}$ is fixed. (B) The height deformation is also dependent on the bilayer thickness $2h_0$. Simulations were performed with flat DLPC membrane with the varied $2h_0$, the helix insertion $c_{0,ins}$ = 0.3 nm$^{-1}$, and $\Delta h_0$= 0.3 nm.

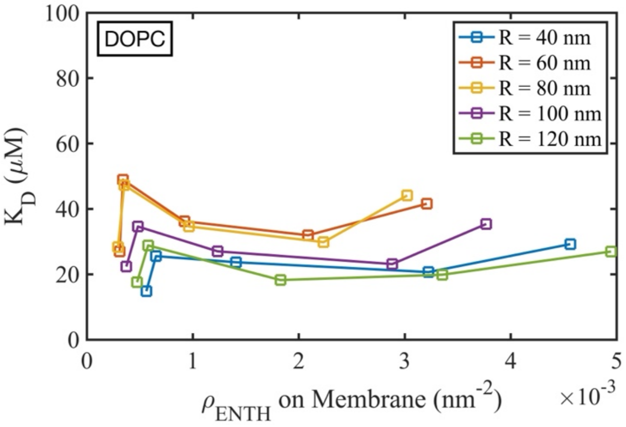


**Figure S5. The dissociation constant ($K_D$) does not exhibit a systematical variation with the ENTH concentration bound to the vesicles.** The experiments were conducted using DOPC lipid vesicles of varying radii ($R$), following the experimental protocol described in the main Methods.
